# T cell engagers control solid tumors through clonal replacement and IL2-driven effector differentiation of CD8 T cells

**DOI:** 10.64898/2025.12.04.692214

**Authors:** Matthias Obenaus, Clara L. Poupault, Christopher S. McGinnis, Céline Prange, Hua Jiang, Leon L. Su, Xiaojing Chen, Zhuang Miao, Joseph J. Muldoon, Winnie Yao, Deepa Waghray, Qinli Sun, Justin Eyquem, Rogelio A. Hernández-López, Ansuman T. Satpathy, Julien Sage, K. Christopher Garcia

## Abstract

Bispecific T cell engagers (TCEs) often exhibit limited efficacy in solid tumors, in part due to immunosuppressive cues in the tumor microenvironment and low expression of targetable tumor antigens. Therapeutic strategies to improve TCE target sensitivity and enhance T cell effector functions therefore have significant translational potential. Here, we engineered TCEs that induce T cell activation *in vitro* against the low-abundance target antigens, TRP2/Kb and DLL3. Despite *in vitro* activity in these models, TCE monotherapy showed limited control of tumor growth in immunocompetent mice. Leveraging this *in vivo* model of TCE treatment failure, we discovered that co-treatment with TCE and a CD25-biased Interleukin-2 (IL2) rescues anti-tumor activity. Further, multimodal single-cell transcriptomic and immune repertoire analyses revealed that TCE-IL2 combination therapy controlled tumors by recruiting and activating new CD8^+^ T cells into the tumor microenvironment. These findings demonstrate that TCE-mediated anti-tumor responses function through a CD8^+^ T cell clonal replacement mechanism that can be augmented by cytokine therapy.

**One Sentence Summary:** Combining TCE therapy with IL-2 enhances TCE efficacy in aggressive small-cell lung cancer and melanoma models with low target antigen density through T cell clonal replacement and CD8^+^ effector T cell differentiation.

## INTRODUCTION

Cancer immunotherapy has significantly improved patient outcomes across a variety of cancer entities, but only a subset of patients benefit long-term (*1, 2*). For this reason, there is a clinical need for improved immunotherapy approaches. Bispecific T cell engagers (TCEs) are engineered antibodies that simultaneously recognize tumor antigens and the T cell receptor (TCR) complex. TCEs can redirect patient T cells to target and eliminate cancer cells – an approach that has been successful in treating B cell malignancies (*3, 4*). However, overall response rates in solid tumor indications are comparatively low and complete responses are rare (*2, 5, 6*). Potential reasons underlying poor TCE responses in solid tumors include an immunosuppressive tumor microenvironment (TME), toxicity due to on-target off-tumor activity, and low expression of targetable tumor antigens (*7*). Optimized TCE treatment regimens that account for these unique challenges have not been thoroughly studied.

Although there are abundant preclinical and mechanistic data on several other immunotherapeutic approaches such as checkpoint blockade (e.g., anti-PD-1 or anti-PD-L1) and adoptive cell therapies (*8*), analogous work on TCEs is comparatively sparse. Indeed, most preclinical studies of TCEs have focused on highly abundant proof-of-concept antigens (*9, 10*), and/or have used mouse xenograft systems that do not model the TME (*11–13*). Therefore, our understanding of how TCEs promote anti-tumor responses in the context of a fully intact immune system, including TCE therapy failure mechanisms, remains incomplete.

In this study, we developed TCEs capable of recognizing low-abundance target antigens on the surface of solid tumor cells. We specifically focused on tumor models that typically respond poorly to checkpoint blockade, including the B16F10 melanoma and small cell lung cancer (SCLC) models (*14–17*). Notably, SCLC tumors respond poorly to clinical checkpoint-inhibitor monotherapy despite a high tumor mutational burden (*18*). Moreover TCE therapy for SCLC is highly relevant given the recent approval of Tarlatamab, a human DLL3 TCE, as a second-line treatment for patients with extensive-stage SCLC (*2, 19*).

We found that low target antigen density limited TCE effectiveness *in vivo*, which can be overcome by combination therapy with a CD25-biased variant of the cytokine IL2 (Interleukin-2). Using this model, we further investigated the mechanistic basis for TCE treatment failure under conditions of low antigen density using multimodal single-cell genomics. This analysis revealed that TCE-mediated *in vivo* tumor control was associated with the recruitment of CD8^+^ tumor-infiltrating lymphocytes (TILs) from the periphery (i.e., clonal replacement; *20*) and induction of anti-tumor effector responses. Overall, our results provide a mechanistic framework that can inform future TCE-based treatment regimens.

## RESULTS

### Optimal geometry is critical for TCE-mediated T cell activation against targets with low antigen expression

To interrogate mechanisms of TCE dysfunction, we developed a model system to test TCE efficacy tuning across a range of target antigen densities (**Fig. 1A**). We first focused on the melanoma model antigen tyrosinase-related protein 2 (TRP2) that is naturally presented at very low levels by the mouse major histocompatibility complex class I (MHC-I) antigen H2-K^b^ on the surface of B16F10 melanoma tumor cells (*21*). Specifically, TRP2/K^b^ was undetectable by flow cytometry, but could be detected by single-molecule imaging at ∼ 44 target molecules per cell (ΔMFI < 100; *21*) To model intermediate antigen densities, we modulated TRP2/K^b^ abundance through Interferon γ treatment (IFNγ; ΔMFI: 474) and TRP2 peptide pulsing of B16F10 cells (ΔMFI 229; **Fig. 1A**). We combined both IFNγ treatment and TRP2 peptide pulsing to model high antigen densities (ΔMFI: 12,724; **Fig. 1A**). Lastly, we established MC38 colon cancer lines transduced to express TRP2 at intermediate to high levels (MC38^low^, MC38^int^, MC38^hi^; ΔMFI: 365-3,847), enabling analyses across a large range of antigen densities.

**Figure 1:**
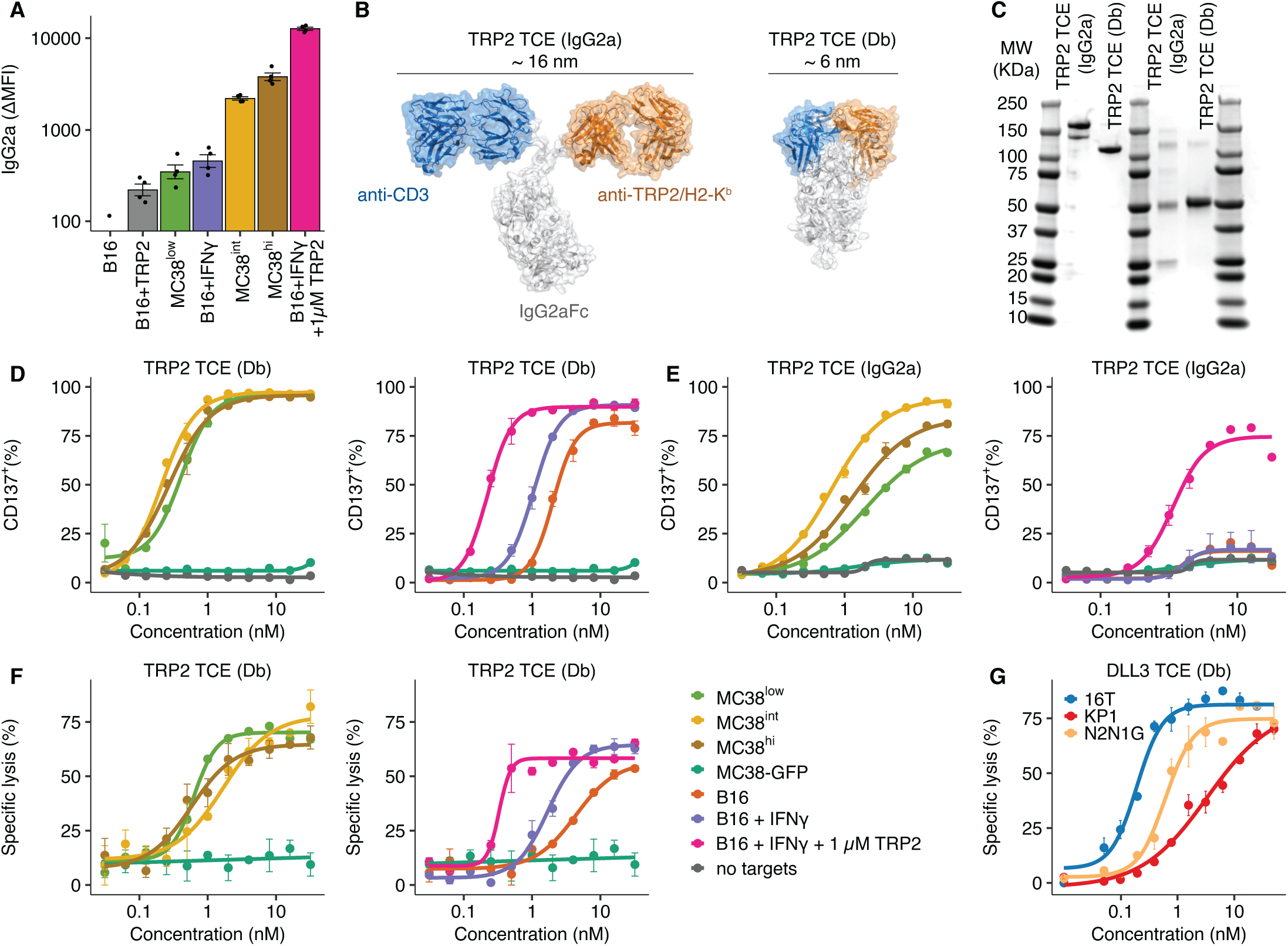
Compact geometry improves TCE activity against targets with low antigen density *in vitro*. (A) TRP2/H2-K^b^ antigen expression on different tumor cell lines: B16F10 were either left untreated or incubated with 100 ng/mL IFNγ and/or 1 µM TRP2 peptide (SVYDFFVWL) for 24 hours and stained with 100 nM TRP2 TCE or no antibody followed by secondary staining with an anti-mouse IgG2a antibody. Shown is the difference between TRP2 TCE-stained and secondary antibody only for each cell line and condition as indicated. The same staining was performed for MC38 cells transduced to stably express different levels TRP2. (B) Representation of the designed TCE variants: a full-length IgG2a-based TCE (left) with the retained flexible hinge domain, and a compact diabody TCE (right), also including the CH2 and CH3 domains of IgG2a. The approximate distances between the CD3 binding paratope and target binding paratope are indicated. To prevent Fc-mediated effector functions, all TCEs carried the D265A mutation. (C) Polyacrylamide gel-electrophoresis of TRP2 Db and full-length IgG2a TCE variants under non-reducing (left) and reducing conditions (right). (D) Effector T cells from C57BL/6 mice were co-cultured with the indicated target cells (Effector-to-target ratio, 1:1) for 36 hours in the presence of TRP2 diabody TCE at the indicated concentration. B16F10 cells were pretreated with IFNγ and/or TRP2 peptide as described in (A) prior to T cell addition. T cell activation was determined by the fraction of CD8^+^ T cells expressing CD137. (E) T cell activation as measured by CD137 upregulation in the presence of TRP2 TCE (full length IgG2a). Co-culture conditions are identical to those described in (D). (F) Cytotoxicity (specific lysis) of OT-I effector T cells (Effector-to-target ratio, 1:1) against the indicated cell lines was measured after 36 h of coculture. B16F10 cells were pretreated with IFNγ and/or TRP2 peptide as described in (A) prior to T cell addition. (G) Cytotoxicity (specific lysis) after 36 h induced by DLL3 TCE against murine SCLC cell lines with different antigen expression levels. All values represent mean ± s.e.m. of intra-assay duplicates unless otherwise specified. In D and F one outlier was removed for cell lines MC38^low^ and MC38^hi^ at a 16 nM concentration due to pipetting error. See also Supplemental Figure S1.

Different strategies to generate TCEs result in a variety of molecular geometries (*22*), prompting us to investigate how TCE geometry affects antigen sensitivity. Because the distance between T cell and target cell membrane is a known factor influencing the efficiency of T cell activation (*23, 24*), we designed full-length IgG2a (hereafter: IgG2a) and compact diabody (Db) TCE variants that result in distinct distances between target- and CD3-binding sites (IgG2a: ∼16 nm; Db: ∼6 nm; **Fig. 1B & C**, **Supplemental Fig. S1A**). Notably, we used a T cell receptor (TCR)-like antibody targeting TRP2/K^b^ that closely resembles the binding geometry of natural TCRs, building on previous work (*21*).

The Db TCE induced T cell activation, as measured by upregulation of the costimulatory marker CD137, with higher sensitivity than the IgG2a TCE (**Fig. 1D & E**, **Supplemental Fig. S1B**). Importantly, only the Db TCE was able to activate T cells and induce cytotoxicity in the presence of untreated B16F10 target cells with the lowest antigen densities (**Fig. 1D-F**, orange line with B16 cells, **Supplemental Fig. S1C & D**). These data illustrate that optimized TCE geometry can facilitate effector-target interactions in low-antigen contexts.

Building on this framework, we additionally engineered mouse Db TCEs recognizing the clinically established small cell lung cancer (SCLC) target antigen DLL3. Specifically, we tested Db TCE variants using three different DLL3 binding domains (**Supplemental Fig. S1E & F**; *25*, 26). The most potent DLL3 Db TCE (based on the DLL3-4 binder) was chosen for further experiments and similarly induced lysis of murine SCLC cell lines with a range of target antigen densities (16T: DLL3^hi^ (antibody bound per cell, ABC: 750); N2N1G: DLL3^int^ (ABC: 377); KP1: DLL3^low^ (ABC: 200)) (**Fig. 1G**, **Supplemental Fig. S1G & H**). Due to their higher sensitivity and potency, all further experiments were performed with Db TCEs.

### TCE monotherapy is effective when antigen is abundant

We next evaluated whether the TCE activity against low-abundance targets observed *in vitro* was recapitulated in syngeneic, immunocompetent mouse models. Specifically, we subcutaneously injected tumor cells expressing DLL3 or TRP2/K^b^ at varying levels and measured tumor volume over time (**Fig. 2A**). For the TRP2/K^b^ antigen, we used an MC38 clone overexpressing TRP2 (MC38^int^) as an example of high antigen density and unmodified B16F10 as an example of low antigen density. For the DLL3 antigen, we selected 16T cells expressing high levels of DLL3, and KP1 cells expressing low levels of DLL3. MC38^int^ tumors were cleared in TRP2 TCE-treated mice (**Fig. 2B-C**, **Supplemental Fig. S2A**), while B16F10 tumors exhibited a non-significant dose-dependent delay in tumor outgrowth and a corresponding increase in tumor T cell infiltration (**Fig. 2D-E, Supplemental Fig. S2B-D**). These results mirrored our *in vitro* data in the same B16F10 and MC38-TRP2 models, where TCE-induced T cell activation and target cell killing correlated with antigen density. Tumor clearance was also observed in DLL3 TCE-treated mice bearing 16T SCLC tumors (high DLL3 density, **Fig. 2F & G**, **Supplemental Fig. S2E**), while KP1 SCLC tumors (low DLL3 density) exhibited non-significant growth delays and increased T cell infiltration but were not cleared (**Fig. 2H & I, Supplemental Fig. S2F**).

**Figure 2:**
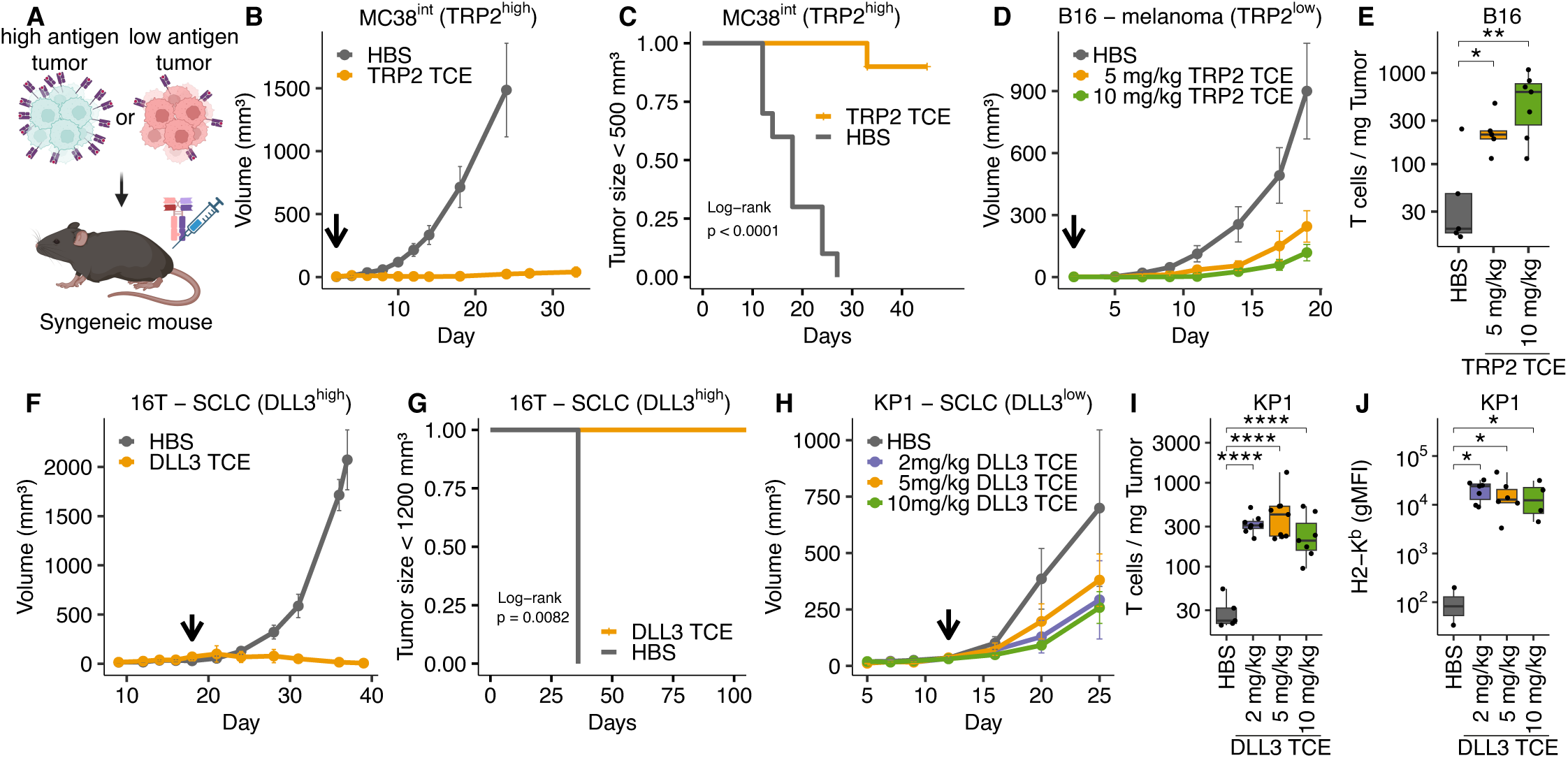
TCE monotherapy rejects tumors with high antigen density but fails to control those with low antigen density. (A) Experimental overview. C57BL/6 mice were injected subcutaneously (s.c.) with either high antigen density or low antigen density tumor cell lines followed by TCE monotherapy. (B) 1 x 10^6^ MC38^int^ cells transduced to express high levels of TRP2 were grafted s.c. onto C57BL/6 mice in Matrigel, TRP2 TCE monotherapy (10 mg/kg q2d) or vehicle control (HBS) was started on day 2 (arrow) after tumor grafting. (p<0.0001, linear mixed-effect modeling) (n=10 mice per group). (C) Fraction of mice with tumors < 500 mm³ during treatment with TRP2 TCE or control (HBS). (D) 0.5 x 10^6^ B16F10 (TRP2 low) were injected s.c. into C57BL/6 mice, TRP2 TCE monotherapy (q2d) or vehicle control (HBS) at the indicated doses was started on day 2 after tumor grafting (HBS: n=6; 5 mg/kg TRP2 TCE: n=6; 10 mg/kg TRP2 TCE n=7 mice). Arrow indicates treatment start. Tumor growth rates were not significantly different between groups (linear mixed-effect modeling). (E) Number of tumor-infiltrating T cells / mg tumor in tumors from mice treated in (D). (F) 0.5 x 10^6^ 16T SCLC cells (DLL3 high) were injected s.c. into B6129SF1/J mice in Matrigel and treated with 5 mg/kg DLL3 TCE or HBS q4d, arrow indicates treatment start (p = 0.0012, linear mixed-effect modeling; HBS: n=3 mice; DLL3 TCE: n=5 mice). (G) Fraction of mice treated in (F) with tumor sizes < 1200 mm³. (H) 0.5 x 10^6^ KP1 SCLC cells (DLL3 low) were injected s.c. into B6129SF1/J mice in Matrigel and treated with the indicated doses of DLL3 TCE or HBS q4d, arrow indicates treatment start. Tumor growth rates were not significantly different between groups (linear mixed-effect modeling; HBS: n=5; 2 mg/kg DLL3 TCE: n=8; 5 mg/kg DLL3 TCE: n=10; 10 mg/kg DLL3 TCE: n=8 mice). (I) Number of tumor-infiltrating T cells / mg tumor in tumors from mice treated in (H). (J) Expression of H2-K^b^ on tumor cells from control or DLL3 TCE-treated mice was measured by flow cytometry as an indicator of TCE-induced IFNγ secretion by T cells. Comparison of means in (E), (I), (J): Student’s t-test; *: p ≤ 0.05, **: p ≤ 0.01, ***: p ≤ 0.001, ****: p ≤ 0.0001. See also Supplemental Figure S2.

Next, we looked for evidence of TIL activation beyond the observed increased T cell numbers. KP1 tumors express very low amounts of MHC-I, which can be upregulated by IFNγ treatment (**Supplemental Fig. S2G**). We measured H2-K^b^ surface expression on KP1 cells as a proxy for TCE-induced IFNγ secretion in tumors and found elevated H2-K^b^ surface abundance after TCE treatment compared to controls (**Fig. 2J**). In summary, these results demonstrate the efficacy of TCE monotherapy against tumors with high antigen density. However, TCE efficacy was limited in low antigen density conditions, despite the potent *in vitro* activity and *in vivo* evidence of TCE-mediated antigen-specific TIL activation.

### Addition of IL2 improves TCE activity

We next sought to define how TCE-mediated control of low antigen density tumors was compromised *in vivo*. T cells have distinct activation thresholds for different effector functions (*27, 28*). For example, cytokine secretion and proliferation are known to require higher degrees of signaling strength compared to cytotoxicity (*29*). We hypothesized that TCE-mediated T cell activation against low antigen density tumors was sub-optimal, leading to limited anti-tumor effects. To test this, we measured IFNγ and IL2 secretion in response to TRP2 TCE-driven T cell activation at low antigen densities. Specifically, we pulsed OT-I splenocytes with low amounts (0.13 nM) of either the ovalbumin-derived peptide (SIINFEKL, Ova N4) or TRP2 peptide (SVYDFFVWL) in the presence or absence of the TRP2/K^b^ TCE (**Fig. 3A**). Since Ova N4 is a strong agonist for the OT-I TCR and is also presented by H2-Kb, it served as a positive control for an ideal T cell response (*30*).

**Figure 3:**
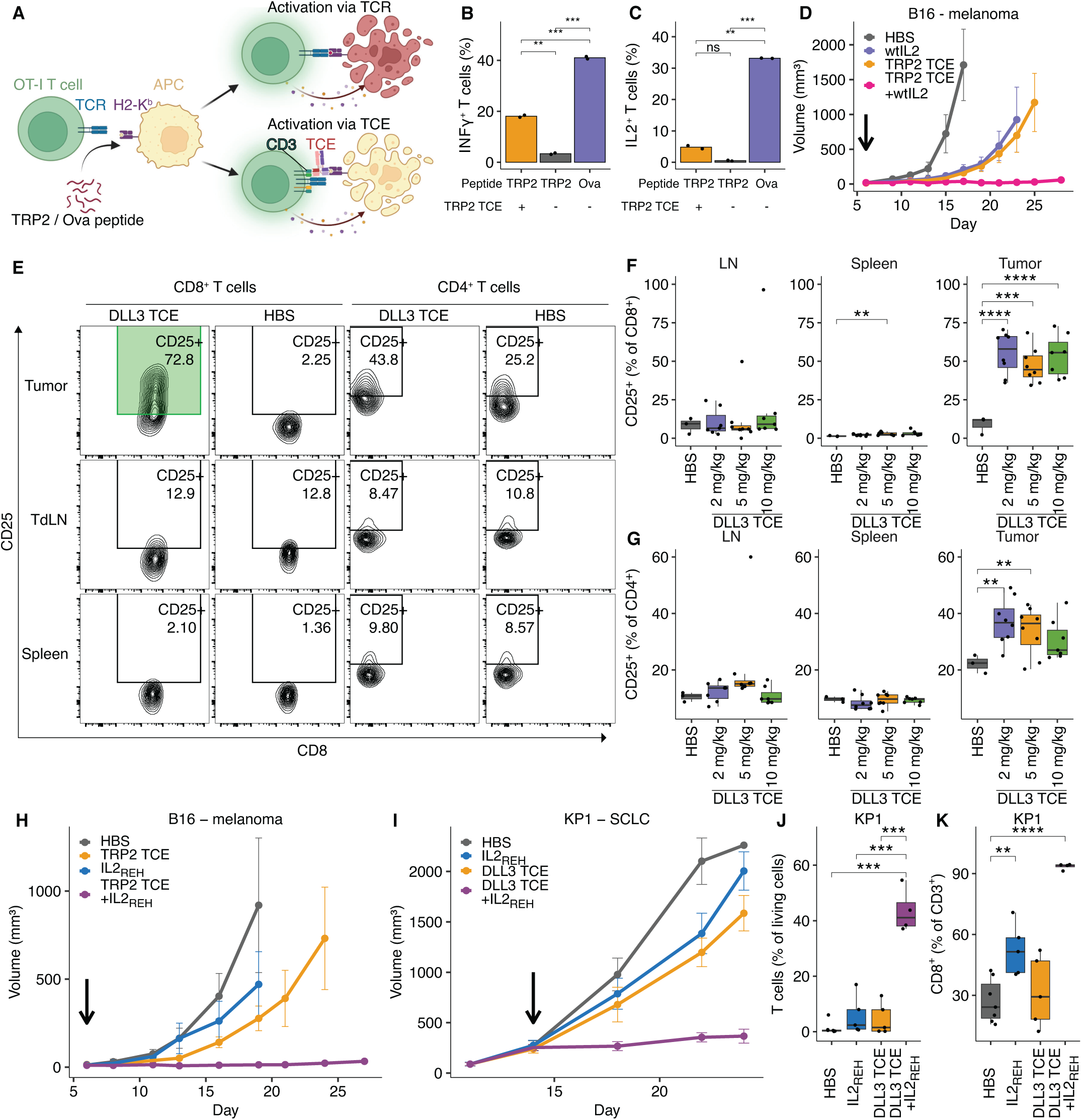
Supplementation with IL2 restores TCE activity in tumors with low antigen density. (A) Procedure to determine T cell cytokine responses after activation via the TCR or TCE. OT-I splenocytes were pulsed with low amounts (0.13 nM) of either Ova N4, resulting in stimulation via the TCR, or TRP2 peptide, resulting in no stimulation (without TRP2 TCE) or stimulation via the TCE (in the presence of 1 nM TRP2 TCE). (B) OT-I splenocytes were prepared as described in (A) and stimulated for 12 h, followed by fixation and intracellular cytokine staining. Bar graphs show the fraction of OT-I T cells positive for IFNγ. (C) Fraction of IL2-positive OT-I T cells. Values depict means of intra-assay duplicates, means-comparison: Student‘s t-test; *: p ≤ 0.05, **: p ≤ 0.01, ***: p ≤ 0.001, ****: p ≤ 0.0001. (D) 0.5 x 10^6^ B16F10 (TRP2 low) were injected s.c. into C57BL/6 mice and treated with either HBS, TRP2 TCE (5 mg/kg, q6d) or murine IL2 (3 µg, q2d) monotherapy, or the combination of TCE and IL2. Arrow indicates treatment start. Linear mixed-effects model, pairwise comparisons: IL2 vs. TCE + IL2: p<0.0001, TCE vs. TCE + IL2: p<0.0001, HBS vs. TCE + IL2: p<0.0001, HBS vs. IL2: p < 0.05, HBS vs. TCE: p < 0.05, p-adjustment by Holm-method (HBS: n=4, IL2: n=3, TCE: n=6, TCE + IL2: n=6 mice). (E) Representative flow cytometry plots showing IL2Rα/CD25 upregulation on tumor-infiltrating T cells during TCE treatment. T cells were isolated from tumors, tumor-draining lymph nodes (TdLN), or spleens from mice harboring KP1 SCLC tumors under DLL3 TCE monotherapy or HBS treatment. CD25 upregulation was most pronounced on CD8^+^ TILs (highlighted in green). (F) Fraction of CD25-expressing CD8^+^ T cells in TdLNs, spleens, and tumors under DLL3 TCE treatment. (G) Fraction of CD25-expressing CD4^+^ T cells in TdLNs, spleens, and tumors under DLL3 TCE treatment. (H) 0.5 x 10^6^ B16F10 (TRP2 low) were injected s.c. into C57BL/6 mice and treated with either TRP2 TCE monotherapy (5 mg/kg, q6d) or the CD25 biased IL2 variant IL2_REH_ (3 µg, q2d), HBS or the combination of TCE and IL2_REH_. Arrow indicates treatment start. Linear mixed-effects model, pairwise comparisons: TCE vs. TCE + IL2_REH_: p<0.0001, HBS vs. TCE: p < 0.05, HBS vs. TCE + IL2_REH_: p<0.0001, IL2_REH_ vs. TCE + IL2_REH_: p = 0.0004, p-adjustment by Holm-method (HBS: n=5, IL2_REH_: n=5, TCE: n=11, TCE + IL2_REH_: n=11 mice). (I) 0.5 x 10^6^ KP1 SCLC cells (DLL3 low) were injected s.c. into B6129SF1/J mice and treated with either DLL3 TCE monotherapy (5 mg/kg, q4d), IL2_REH_ (3 µg, q2d), HBS or the combination of TCE and IL2_REH_. Arrow indicates treatment start. Linear mixed-effects model, pairwise comparisons: TCE vs. TCE + IL2_REH_: p<0.05, IL2_REH_ vs. TCE + IL2_REH_: p <0.0001, HBS vs. TCE: p <0.0001, TCE vs. TCE: p <0.0001, HBS vs. TCE + IL2_REH_: p <0.0001, p-adjustment by Holm-method (n= 10 mice per group). (J) Fraction of αβ T cells as a fraction of CD45^+^ hematopoietic cells in tumors from mice treated in (I). (K) Fraction of CD8^+^ T cells as a fraction of all αβ T cells in tumors from mice treated in (I) Comparison of means in (F), (G), (J), (K): Student’s t-test; *: p ≤ 0.05, **: p ≤ 0.01, ***: p ≤ 0.001, ****: p ≤ 0.0001. See also Supplemental Figures S3 and S4.

Ova N4 pulsed OT-I splenocytes, receiving stimulation via their natural ligand (i.e., Ova N4), secreted IFNγ and IL2 (**Fig. 3B & C**). In contrast, TRP2-pulsed splenocytes receiving stimulation via the TCE exhibited lower IFNγ and almost no IL2 secretion (**Fig. 3B & C**). These results are in line with previous studies showing that TCEs are poor inducers of cytokine secretion – for example, the clinically used DLL3 TCE Tarlatamab induces robust cytotoxicity but only minimally stimulates cytokine secretion (*31*).

IL2 has multiple roles in the regulation of immune responses, including during T cell priming and stimulation of T cell effector functions (*32*). Since IL2 secretion was particularly limited in response to TCE stimulation, we reasoned that the addition of recombinant IL2 to TCE treatment would restore anti-tumor activity in contexts of low tumor antigen density. After identifying a tolerable dose of IL2 in combination with TCE (3 µg IL2 every 2 days + 5 mg/kg TCE**; Supplemental Fig S3A**), we treated B16F10 tumor-bearing mice with vehicle, TCE or IL2 monotherapy, or TCE-IL2 in combination and observed enhanced tumor control specifically after combination therapy (**Fig. 3D, Supplemental Fig. S3B**).

Because of the known clinical side effects associated with recombinant IL2 treatment (*33*), we next explored strategies to target the anti-tumor IL2-effect to the TME. We noted that CD25/IL2Rα was predominantly upregulated by KP1 CD8^+^ TILs but not splenic or tumor-draining lymph node T cells after treatment with the DLL3 TCE (**Fig. 3E**-**G**). This observation led us to test an IL2 variant (IL2_REH_) with lower affinity towards the common gamma chain (γ_c_) and consequently stronger dependence on CD25 (*34*). Importantly, the human version of CD25-targeted IL2 is currently being investigated as monotherapy in melanoma patients (*35*). As was observed with wild-type IL2, the TCE-IL2_REH_ combination controlled the growth of both B16F10 and KP1 tumors (**Fig. 3H & I**, **Supplemental Fig. S3C & D**). Flow cytometry analysis of KP1 tumors from the different treatment groups showed that combination therapy further increased the TIL fraction compared to IL2_REH_ or TCE monotherapy (**Fig. 3J**). Notably, the T cell infiltrate was dominated by CD8^+^ TILs in the combination therapy group (**Fig. 3K**). Overall, TCE-IL2_REH_ combination therapy was well tolerated without overt signs of acute toxicity (**Supplemental Fig. S4A-C**). Interestingly, surviving mice treated with TRP2 TCE-IL2 (wild-type or IL2_REH_) combination therapy but not TCE monotherapy developed vitiligo and leukotrichia as manifestations of on-target/off-tumor toxicity (**Supplemental Fig. S4D & E**). This observation suggests that the inefficient T cell activation observed during TCE monotherapy is not limited to the TME, but a systemic phenomenon.

### TCE-IL2_REH_ combination therapy promotes anti-tumor TIL responses and differentially affects CD4^+^ and CD8^+^ TILs

Having identified the ability of IL2 to correct TCE dysfunction in low tumor antigen contexts, we used joint single-cell RNA sequencing (scRNA-seq) and TCR sequencing (scTCR-seq) to deeply characterize TIL responses during mono- and combination therapy. Specifically, we treated B16F10 tumor-bearing mice with vehicle, TCE or IL2_REH_ monotherapy, or TCE-IL2_REH_ in combination, and isolated TILs from each group on day 6 after treatment (**Fig. 4A**). MULTI-seq (*36*) was used to barcode pooled TILs from each treatment group (n = 7-8 tumors per group) prior to multiplexed scRNA-seq and TCR-seq analysis using the 10x Genomics system. Following quality-control filtering and MULTI-seq demultiplexing, we used unsupervised clustering and differential gene expression (DGE) analysis to identify five subsets of CD4^+^ TILs and six subsets of CD8^+^ TILs: CD4^+^ naïve (*Tcf7*, *Lef1*), memory (*Zfp36l2*), activated (*Ifng*, *Nr4a2*), regulatory T cells (Treg; *Foxp3*, *Ikzf2*), and proliferative (*Stmn1*) TILs; CD8^+^ naïve (*Sell*, *Tcf7*), effector (*Ly6c2*, *Ctla2a*), progenitor exhausted-like (Tpex-like; *Xcl1*, *Ccr7*), terminally exhausted (Tex; *Pdcd1*, *Havcr2*), proliferative exhausted (Tex-prolif; *Pdcd1*, *Ccna2*), and interferon-stimulated (IFN; *Isg15*, *Ifit3*) TILs (**Fig. 4B; Supplemental Fig. S5A**).

**Figure 4:**
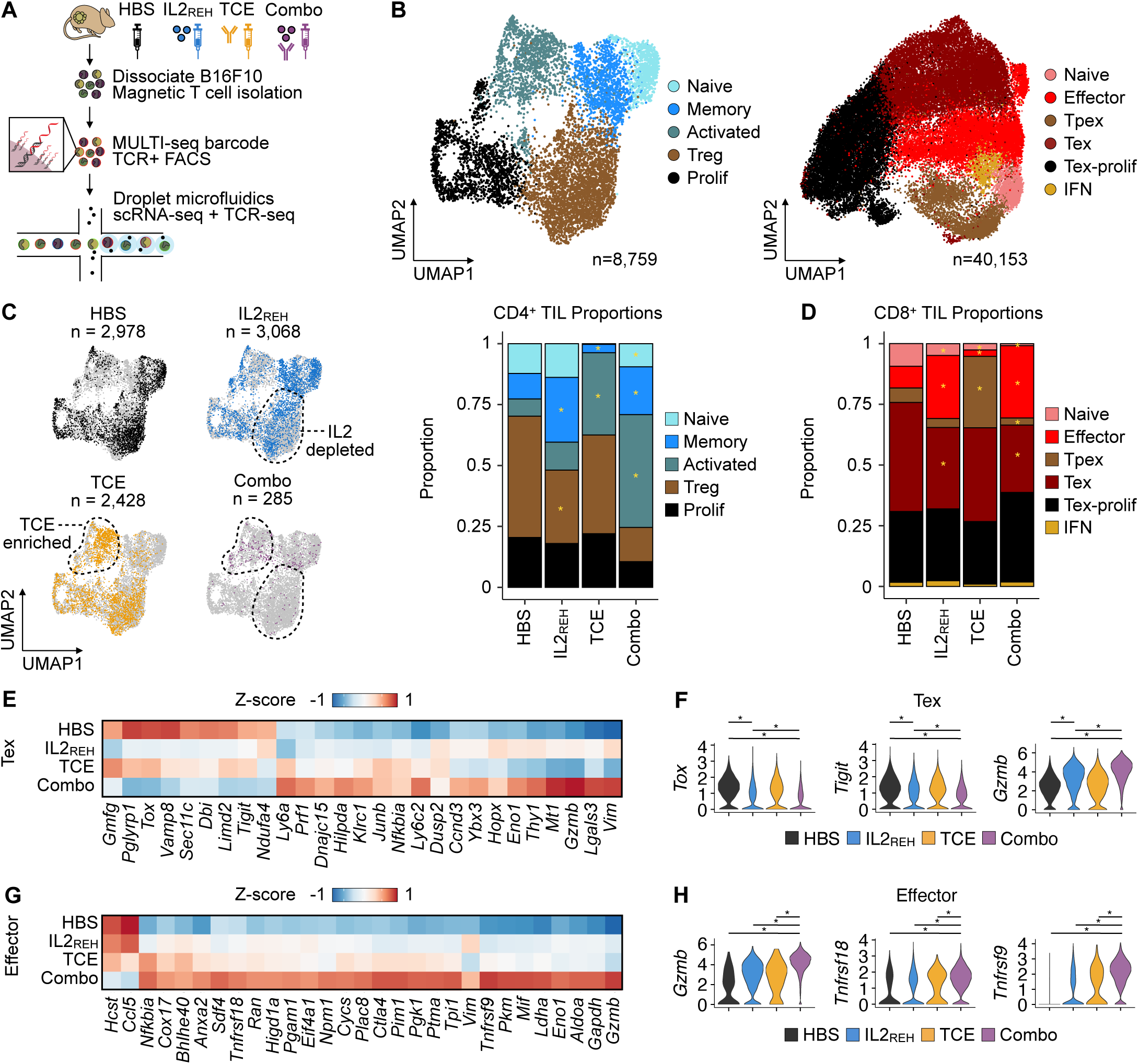
TCE-IL2_REH_ combination therapy promotes anti-tumor TIL responses and differentially affects CD4^+^ and CD8^+^ TILs. (A) Schematic overview of experimental workflow. Tumors were removed at day 6 after treatment prior to MULTI-seq barcoding, T cell enrichment, and paired scRNA-seq and scTCR-seq. (B) Uniform manifold approximation and projection (UMAP) visualization of CD4^+^ TIL (left) and CD8^+^ TIL (right) gene expression space colored by subtype annotation. (C) UMAP visualization of CD4^+^ TIL gene expression space colored by treatment groups (left) with bar charts of CD4^+^ TIL subtype proportions in each treatment group. * p < 0.05 and proportion ratio ± 0.5, propeller test. (D) Bar charts of CD8^+^ TIL subtype proportions in each treatment group. * p < 0.05 and proportion ratio ± 0.5, propeller test. (E) Z-score heatmap of the average expression of treatment-specific Tex marker genes in each treatment group. Genes grouped by hierarchical clustering of the z-score matrix. (F) Violin plots of a subset of treatment-specific Tex marker genes in each treatment group. * p < 0.05 Bonferroni-corrected Wilcoxon rank-sum test. (G) Z-score heatmap of the average expression of treatment-specific effector CD8^+^ TIL marker genes in each treatment group. Genes grouped by hierarchical clustering of the z-score matrix. (H) Violin plots of a subset of treatment-specific effector CD8^+^ TIL marker genes in each treatment group. * p < 0.05 Bonferroni-corrected Wilcoxon rank-sum test. See also Supplemental Figure S5.

Analyses of CD4^+^ TIL subtype proportions in each condition revealed variable signatures indicative of immunogenic tumor control in each treatment group. Specifically, IL2_REH_ therapy was associated with decreased Treg frequencies compared to vehicle-treated controls – mirroring prior reports that CD25-biased IL2 increases the CD8^+^ TIL:Treg ratio in the tumor microenvironment (**Fig. 4C**; *33, 37*). Since Tregs have established immunosuppressive functions in the TME (*38*), the decrease in Tregs driven by IL2_REH_ monotherapy provides a possible explanation for the observed modest tumor growth delay induced by IL2_REH_ monotherapy. In contrast, TCE therapy was linked to increased frequencies of activated CD4^+^ TILs (**Fig. 4C**), indicative of productive TCR binding by the TCE despite eventual tumor outgrowth. TCE-IL2_REH_ combination therapy reflected an additive immunomodulatory effect of IL2_REH_ (i.e., decreased Treg frequency) and TCE (i.e., increased activation) and was associated with an increased CD8^+^:CD4^+^ TIL ratio (**Supplemental Fig. S5B**), recapitulating the flow cytometry data (**Fig. 3K**).

Changes in CD8^+^ TIL subtype proportions across treatment groups revealed a deviation from the additive responses observed in CD4^+^ TILs. For example, TCE monotherapy was predominantly associated with an increase in Tpex-like CD8^+^ TILs relative to all treatment groups, which is consistent with productive TCR engagement and subsequent CD8^+^ TIL activation, but incomplete transition to a full effector phenotype (**Fig. 4D**). In contrast, both IL2_REH_ treatment groups were associated with increased CD8^+^ effector TILs and a reduction in Tex cells. Notably, the IL2_REH_-induced changes in CD8^+^ effector and Tex proportions were more pronounced following combination therapy (**Supplemental Fig. S5C**), suggesting that TCE-IL2_REH_ amplified this response.

Indeed, DGE analysis of Tex cells in each treatment group revealed an IL2_REH_-specific transcriptional signature that was amplified after combination therapy (**Fig. 4E**). This signature included downregulated T cell exhaustion markers (*Tigit*, *Tox*) and upregulation of the T cell cytotoxicity marker *Gzmb* (**Fig. 4F**), suggesting an increased ability of CD8^+^ TILs to promote anti-tumor responses following combination therapy. In contrast, DGE analysis of activated CD8^+^ TILs revealed a transcriptional signature induced specifically by combination therapy that included many glycolytic enzymes (**Fig. 4G; Supplemental Fig. S5D)** – a known metabolic shift associated with effector T cell function (*39–41*) – as well as genes linked to T cell activation and effector function (*Gzmb*, *Tnfrsf18*/*4-1BB*, *Tnfrsf9*/*GITR*; **Fig. 4H**). Collectively, these results illustrate how combination therapy altered TIL population structure (e.g., reduction of Tregs, activation of CD4^+^ TILs, and induction of effector CD8^+^ TILs) and transcriptional state (e.g., differentially-expressed CD8^+^ TIL exhaustion, cytotoxicity, and metabolic markers) in ways that are consistent with immunogenic anti-tumor control.

### TCE-mediated tumor growth control is associated with clonal replacement followed by activation of CD8^+^ TILs

T cell immunotherapy-mediated anti-tumor responses are driven by two distinct mechanisms – stimulation of existing tumor-specific T cell clones in the TME (i.e., clonal expansion) or recruitment and activation of novel peripheral T cell clones into the TME (i.e., clonal replacement). scTCR-seq analysis can measure the clonal relationships between T cells in the TME and thereby help distinguish these two models. For example, scTCR-seq was leveraged to demonstrate that clonal replacement was the dominant mechanism driving anti-PD-1 therapy responses in basal or squamous cell carcinoma patients (*20*). However, unlike during checkpoint blockade, TCEs indiscriminately bind to and activate both tumor antigen-specific and non-specific T cells, raising the question of which mechanism dominates the TCE-induced anti-tumor effect.

We analyzed scTCR-seq data across treatment groups (vehicle, TCE, IL2_REH_, or TCE-IL2_REH_ combination) from a single time-point to assess the relative contributions of clonal replacement and expansion during TCE-mediated anti-tumor responses. We hypothesized that although both mechanisms would result in increased overall TIL numbers, anti-tumor responses driven by clonal replacement would be associated with treatment-specific increases in clonal diversity.

Global analysis of clone frequencies across treatment groups revealed that both TCE monotherapy and combination therapy were associated with increased clonal diversity relative to IL2_REH_ monotherapy or vehicle controls (**Fig. 5A**). Specifically, among CD8^+^ TILs with detectable TCR sequences, 58.2% (3,889/6,682 total cells) and 65.4% (6,542/10,001) of CD8^+^ TILs were associated with unique clones in the TCE monotherapy and combination therapy groups, respectively. In contrast, only 32.2% (1,014/3,143 total cells) and 39.4% (4,180/10,605) of CD8^+^ TILs were respectively associated with unique clones in the vehicle control and IL2_REH_ monotherapy groups. Further, analysis of subtype biases among clones revealed minimal clonal overlap between exhausted and non-exhausted subtypes (**Fig. 5B**); and notably, Tex and Tex-prolif CD8^+^ TILs were enriched for expanded clones compared to the polyclonal naive- and Tpex-like subsets (**Fig. 5B**, top; **Fig. 5C**). This enrichment for high-abundance clones within exhausted CD8^+^ TILs is consistent with the expected expansion of pre-existing tumor antigen-specific T cells at baseline, as has been observed in human patients (*20*). Considered collectively, the overall increase in clonal diversity after TCE monotherapy and combination therapy, along with the previously-discussed unique association between TCE monotherapy and clonally-diverse Tpex-like CD8^+^ TILs (**Fig. 4D; Fig. 5C**), are consistent with TCE-mediated activation and recruitment of polyclonal T cells from the periphery.

**Figure 5:**
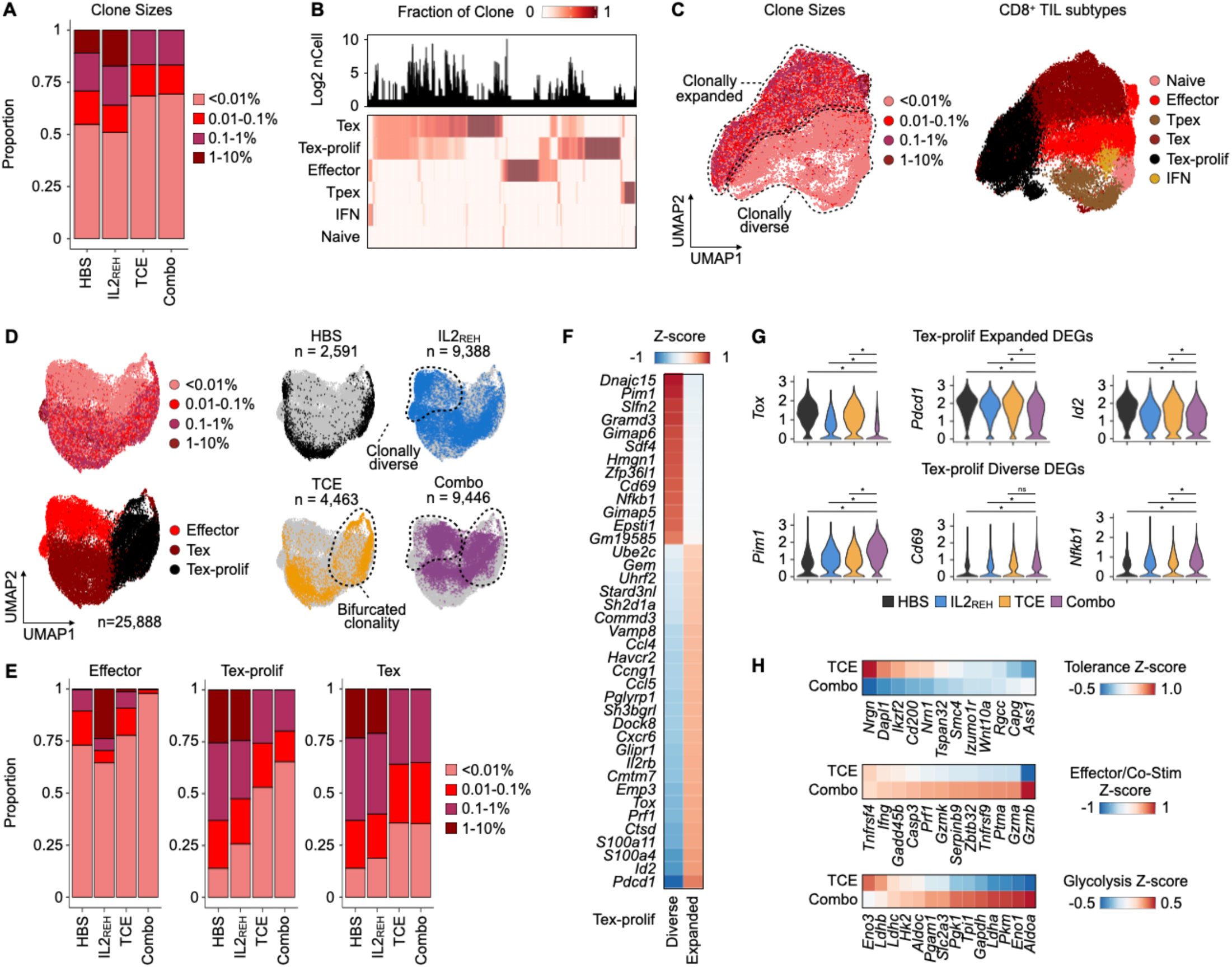
TCE-mediated tumor growth control is associated with clonal replacement followed by activation of CD8^+^ TILs. (A) Bar charts describing the proportions of rare (<0.01%), small (0.01-0.1%), medium (0.1-1%), and large (1-10%) CD8^+^ TIL clone sizes represented in each treatment arm. (B) Heatmap depicting the phenotypic distribution of individual TCR clones across CD8^+^ TIL subsets (bottom) along with histogram of TCR clone frequency (top). Individual clones are grouped by hierarchical clustering of the clone fraction matrix. (C) UMAP visualization of CD8^+^ TIL gene expression space for cells with detectable TCRs colored by clone size (left) or subtype (right). CD8^+^ TIL subtypes associated with rare or expanded clones are highlighted with dotted lines. (D) UMAP visualization of effector, Tex, and Tex-prolif CD8^+^ TIL gene expression space colored by clone size or subtype (left) or by treatment groups (right). CD8^+^ TIL subtypes associated with clonal diversity or bifurcated clonality are highlighted with dotted lines. (E) Bar charts describing the proportions of rare (<0.01%), small (0.01-0.1%), medium (0.1-1%), and large (1-10%) clone sizes among effector, Tex, and Tex-prolif CD8^+^ TILs. (F) Z-score heatmap of the average expression of marker genes distinguishing the clonally-expanded and diverse Tex-prolif populations. Genes grouped by hierarchical clustering of the z-score matrix. (G) Violin plots of a subset of clonally-expanded and diverse Tex-prolif marker genes in the full Tex-prolif population binned by treatment group. (H) Z-score heatmap of the average expression of glycolysis, effector/co-stimulation, and T cell dysfunction gene markers in clonally diverse CD8^+^ TILs subsetted by therapy group (TCE monotherapy and combination therapy highlighted). * p < 0.05 Bonferroni-corrected Wilcoxon rank-sum test.

Next, to interrogate subtype-specific differences in T cell clonality between treatment groups, we focused on CD8^+^ TIL populations that exhibited some degree of clonal expansion: Tex, Tex-prolif, and CD8^+^ effector TILs. Comparing clone frequencies in each subtype across treatment groups revealed two notable signatures. First, IL2_REH_ monotherapy and combination therapy were both enriched for clonally diverse effector CD8^+^ TILs, but clonal diversity was highest in the combination group (**Fig. 5D; Fig. 5E**, left). Second, TCE monotherapy and combination therapy were both associated with a pattern of bifurcated clonality, manifesting as a treatment-specific population of clonally-diverse Tex-prolif cells alongside the clonally-expanded Tex-prolif population that was observed in all treatment groups (**Fig. 5D; Fig. 5E**, middle). Notably, bifurcated clonality was not observed in Tex, even though the overall diversity remained higher in both TCE treatment groups (**Fig. 5D; Fig. 5E**, right).

Unbiased DGE analysis between the clonally expanded and diverse Tex-prolif populations revealed upregulation of exhaustion markers (e.g., *Tox*, *Id2*, and *Pdcd1*) in the clonally expanded pool that were suppressed after combination therapy (**Fig. 5F & G**). In contrast, the clonally diverse Tex-prolif population exhibited increased expression of markers associated with T cell activation (e.g., *Cd69* and *Nfkb1*) and glycolysis (e.g., *Pim1*) that were specifically elevated after combination therapy (**Fig. 5F & G**). Overall, these observations support a model where the combination of IL2_REH_-induced effector functions and TCE-driven recruitment and activation of peripheral T cells leads to tumor growth control.

To further test this model and understand how IL2_REH_ supports TCE-recruited T cells, we compared the gene expression profiles of all diverse clones after TCE monotherapy and combination therapy. T cells activated by TCE monotherapy expressed many genes associated with T cell dysfunction/tolerance (e.g., *Dapl1*, *Nrgn*, and *Ikzf2*/*Helios*; **Fig. 5H**, top), consistent with their predominantly Tpex-like gene expression pattern. Conversely, rare TCR clonotypes in the combination therapy group induced a transcriptional program associated with effector and co-stimulatory CD8^+^ TIL functions (e.g., *Prf1*, *Gzmb*, and *Tnfrsf9*; **Fig. 5H**, middle) and glycolysis (e.g., *Aldoa*, *Eno1*, and *Pgk1*; **Fig. 5H**, bottom), mirroring our observations in the clonally diverse Tex-prolif population.

Taken together, these multimodal scRNA-seq and scTCR-seq analyses demonstrate that TCE therapy recruited a population of clonally diverse CD8^+^ TILs to the TME. In the context of TCE monotherapy, these newly recruited CD8^+^ TILs were characterized by a gene expression signature indicative of dysfunctional T cell activation, while TCE-IL2_REH_ combination therapy limited exhaustion marker expression and induced the expression of effector molecules. These results support a model (**Fig. 6**) where TCE-mediated anti-tumor responses function through a mechanism of clonal replacement and T cells recruited through TCE treatment acquire full effector functions through supplemental exogenous IL2, when antigen is limited.

**Figure 6:**
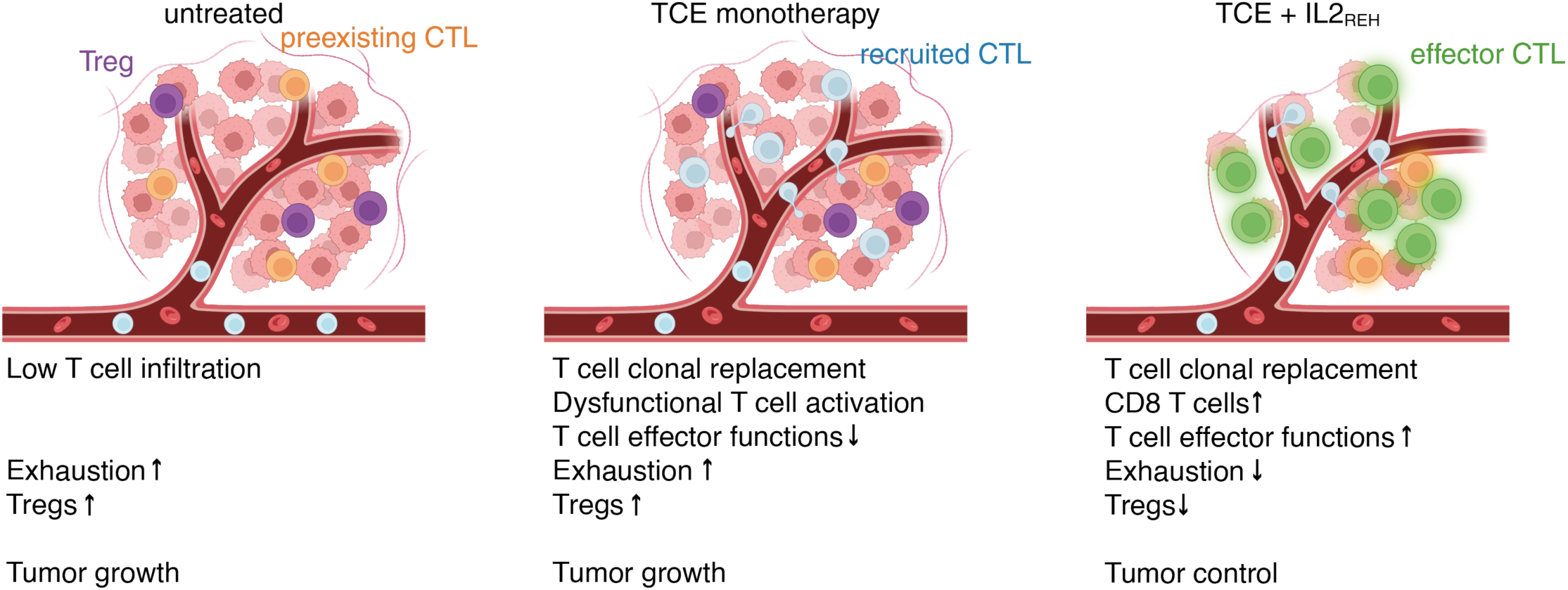
Proposed mechanism underlying TCE treatment failure against tumors with low antigen density and synergy between TCE and IL2_REH_ combination therapy. The untreated TME is characterized by low T cell infiltration and elevated CD8^+^ TIL exhaustion and Treg proportions, leading to unchecked tumor growth. TCE monotherapy drives recruitment of peripheral T cells into the TME which are dysfunctionally activated, leading to increased CD8^+^ TIL clonal diversity without significant changes in exhaustion and cytotoxicity signatures or Treg proportions, resulting in incomplete tumor control. TCE-IL2_REH_ combination therapy recruits peripheral T cells into the TME which are then successfully activated, leading to an increased population of clonally-diverse CD8^+^ TILs that are less exhausted and more capable of anti-tumor activity, resulting in immunogenic tumor control. Figure created using BioRender.

## DISCUSSION

In this study, we used syngeneic melanoma and SCLC models that are characterized by low T cell infiltration, sparse target antigen expression, and poor responses to checkpoint blockade (*14*), to investigate mechanisms of TCE therapy failure.

We first engineered TRP2/K^b^ and DLL3-targeting TCEs that function against tumors with low target antigen density *in vitro*. We found that the compact geometry of diabody TCEs – functioning by reducing the intermembrane distance between T cell and target cell and perhaps more efficient CD45 exclusion (*42, 43*) – improved T cell activation and target cell lysis in response to lower target antigen densities. Next, we used immunocompetent mouse models to assess the *in vivo* efficacy of TCE treatment. Despite the observed *in vitro* killing and *in vivo* evidence of antigen-specific increases in TIL numbers and activation (i.e., MHC-I upregulation on cancer cells and CD25 upregulation on TILs), TCE monotherapy did not control low antigen-density tumor growth, enabling interrogation of TCE therapy failure mechanisms.

We demonstrated that TCE-mediated *in vivo* tumor control was restored through supplemental treatment with IL2_REH_, a CD25-biased IL2 variant. Multimodal single-cell genomic analyses across monotherapy and combination therapy groups revealed the mechanism underlying this growth control – TCE-induced clonal replacement of the CD8^+^ TIL compartment and subsequent IL2_REH_-mediated effector differentiation. In mice treated with TCE alone, newly recruited polyclonal CD8^+^ TILs were biased towards a Tpex-like phenotype, indicating an incomplete T cell activation trajectory that resulted in poor tumor control. In contrast, TCE-IL2_REH_ combination therapy controlled tumors by increasing overall intratumoral T cell numbers and clonal diversity, increasing the CD8:CD4 T cell ratio, reducing the frequency of Tregs within the tumor, and finally, inducing a unique effector gene program reflecting the upregulation of effector, glycolysis and costimulatory markers. Notably, several potential mechanisms may explain the TCE-driven T cell recruitment from the periphery into the TME, including migratory arrest of recirculating T cells (*44*) and/or induction of chemokines (CCL3, CCL4) secreted by preexisting TILs (*45*).

The observed reduction in Tregs in the TME after IL2_REH_ treatment was surprising because Tregs express high amounts of CD25, making them particularly sensitive to IL2_REH_ (*34*). However, our observations are in line with recent reports demonstrating Treg reduction in the TME after treatment with an alternative CD25-biased IL2 variant (N88D; *33*) Importantly, disruption of CD25 binding reduced the anti-tumor effect of IL2 monotherapy in MC38 tumors and the additive effect of IL2 on checkpoint blockade in the context of LCMV infection (*33, 37*). We speculate that IL2_REH_ induction of activated CD8^+^ and/or CD4^+^ TILs outweighs its effect on Tregs within the TME, and that these activated proinflammatory TILs suppress Treg induction.

Notably, our model of successful TCE- and IL2_REH_-mediated tumor control in low-antigen density solid tumors contrasts with recent reports that describe how the number of preexisting memory T cells in samples from multiple myeloma patients predicts response rates to a TCE targeting the B cell maturation antigen (BCMA; *46*). Because our analysis focused on TCE dysfunction in the context of low target-antigen density, further studies are needed to explore whether this difference is related to the higher target-antigen density on myeloma cells, specific differences between multiple myeloma and solid tumor TMEs, or interspecies differences in the T cell response to TCEs.

Overall, our results reveal TCE-induced recruitment of “fresh” CD8^+^ T cells into the TME as a dominant mechanism of TCE-mediated anti-tumor responses, explaining why even tumors with poor prior T cell infiltration can be responsive to TCE therapy. In the context of low target antigen density, TCE-induced T cell activation was dysfunctional until rescued through combination therapy with exogenous IL2. These observations provide a framework guiding the development of clinical TCE combination regimens under these challenging conditions.

### Limitations of the study

Our analysis focused on TCE treatment failure in the context of low-antigen density tumors, identifying dysfunctional T cell activation as the primary failure mechanism, which can be rescued by cotreatment with IL2. However, further studies are needed to determine if TIL gene signatures associated with successful TCE monotherapy against high-antigen tumors resemble those seen with combination therapy or if they are driven by different mechanisms. Second, while we use syngeneic tumor models in immunocompetent mice, allowing us to capture most aspects of the TME, subcutaneous tumor models cannot recapitulate subtle TME variables specific to tumor locations other than the subcutaneous tissue. Lastly, the use of cell lines does not capture the heterogeneity typically present in human cancers.

## MATERIALS AND METHODS

### Design

The goal of this study was to investigate treatment resistance to TCEs in syngeneic cancer models with low tumor target antigen density and poor T cell infiltration, and to develop strategies to enhance TCE responses *in vivo.* To these ends, we tested TCEs *in vitro* on cancer cell lines with varying antigen densities, and *in vivo* by treating immunocompetent mice bearing subcutaneous tumors. C57BL/6 and B6129SF1/J mice were used for the study and were either purchased from the Jackson Laboratory (C57BL/6 and B6129SF1/J) or bred in-house (B6129SF1/J). All mouse experiments were performed in accordance with policies from Stanford University’s Institutional Animal Care and Use Committee (IACUC). At treatment start, mice were randomized into treatment groups (3-11 mice per group) to ensure equal means of tumor volume across groups, and mice with unusually small or large tumors compared to the cohort were excluded before randomization. Mice that died or were euthanized for animal welfare reasons prior to endpoint were excluded from analysis. For mouse studies using the SCLC models, the researcher conducting the tumor measurements was blinded to the treatment groups. Unless specific otherwise in the corresponding Figure legends, no outliers were excluded from analyses.

### Mice

All animal experiments were conducted in accordance with the policies set by Stanford’s institutional animal care and use committee under APLAC protocol numbers 13565 and 32279. For experiments involving B16F10 and MC38, female C57BL/6 mice (4-10 weeks old) were purchased from the Jackson Laboratory. For experiments involving KP1 and 16T cells, B6129SF1/J were either purchased from the Jackson Laboratory or bred in-house.

### Tumor transplants

B16F10 cells were injected subcutaneously at 0.5 x 10^6^ cells per animal in PBS without Matrigel. MC38, KP1, and 16T cells were mixed directly before injection with Matrigel and injected at 1 x 10^6^ (MC38) or 0.5 x 10^6^ (KP1, 16T) cells per animal. Tumors were measured at the indicated timepoints using a caliper. Tumor volume (*V*) was calculated assuming an ellipsoid *V* = 4/3 *π* × *x*/2 × *y*/2 × *z*/2. In cases where tumor height could not be accurately determined and for all measurements in DLL3 and KP1 tumors volume was calculated *V* = 4/3 *π* × *x*/2 × (*y*/2)^2, where *x* ≥ *y*.

### Intraperitoneal treatment administration

For all experiments involving KP1 and 16T tumors, treatment and tumor size measurements were performed by independent personnel and examiners were blinded with respect to treatment groups. Unless otherwise specified, the following dosing schedule was used: 5 mg/kg TRP2 TCE every 6 days, 5 mg/kg DLL3 TCE every 4 days, 3 µg IL2/IL2_REH_ every 2 days. Dose levels of MSA-fused IL2/IL2_REH_ were based on the IL2 component only. On the day of treatment start, animals were randomized to achieve equal means of tumor sizes across groups using the ‘anticlust’ R package (*47*).

### Cell lines and media

The MC38 cell line was purchased from Kerafast. The B16F10 cell line was purchase from ATCC. B16F10-RFP cells were a gift from Anusha Kalbasi. HEK293T, B16F10, MC38 and derived cell lines were maintained in Dulbecco’s modified eagle’s medium (DMEM), supplemented with 10% fetal bovine serum (FBS), L-glutamine (Gibco) and sodium pyruvate (Gibco). SCLC cell lines N2N1G, KP1 and 16T were previously isolated from spontaneous tumors from *Rb1/Trp53* double knockout mice (KP1 and 16T) or lymph node metastasis from *Rb1/Trp53/Rbl2* triple knockout mice (N2N1G; *48, 49*) and were maintained in Roswell Park Memorial Institute Medium (RPMI) 1640 + 10% bovine growth serum + L-Glutamine + Penicillin/Streptomycin. All primary T cells were cultured in RPMI + 10% FBS + 2-ME + sodium pyruvate + MEM non-essential amino acids + L-Glutamine + Penicillin/Streptomycin. Expi293 cells were maintained in Expi293 Expression Medium (Gibco).

### Lentivirus production

3 x 10^6^ HEK293T cells were seeded into 100 mm dishes in 10 mL DMEM + 10% FBS + L-Glutamine + sodium pyruvate. The following day, transfection was performed with FuGeneHD according to the manufacturer’s instructions. Briefly, 2.93 µg psPAX, 1.52 µg pMDM2G, and 4.39 µg gene of interest were diluted into 585 µL Opti-MEM. After addition of 26.5 µL FuGeneHD, complexes were incubated for 15 min. at room temperature. Medium on HEK293T cells was replaced with 10 mL fresh DMEM + 10% FBS + L-Glutamine + sodium pyruvate. Lentiviral particles were harvested after 48 and 72 hours, filtered through a 0.22 µm filter and pooled. Viral particles were concentrated by addition of PEG-8000 to a final concentration of 10% (W/V) and 300 µM NaCl (final concentration) followed by incubation at 4 °C overnight and centrifugation at 1500 g for 30 min at 4 °C. Pelleted virus was resuspended in PBS and added to target cell lines. Protamine sulfate at a final concentration of 4 µg / mL was used during cell transduction.

### Bispecific TCE design

Full length sequences of all antibody constructs are provided in **Supplementary Table S1**. The TRP2/H2-K^b^ specific antibody clone TRP2#13 was previously developed in our lab and was described before (*21*). Clone 2C11 was used for the CD3 binding arm (*50*). Preferential pairing of IgG heavy chain heterodimers was achieved by introducing electrostatically complementary mutations into the CH3 domain of the target-binding and CD3-binding arms, similar to previously described protocols (*51*). Specifically, we introduced K409E and K439D mutations into the CH3 domain of the target binding arm and complementary D399K and E356K mutations into the CH3 domain of the CD3 binding arm. Additionally, the following mutations were introduced into the CH3 domain of both heavy chains: T346S, M368L, T370K, and R411T, as described previously (*51*). A shifted cysteine bridge was used to aid in preferential pairing of light chains with their corresponding heavy chain. Specifically, we removed the original cysteine bridge between heavy and light chain in the CD3-binding arm (C209V in the heavy chain, C214V in the corresponding light chain) and introduced a new cysteine bridge (Y122C on the heavy chain, S121C on the corresponding light chain). The D265A mutation was introduced to abrogate Fc mediated effector functions (*52*). For the diabody format the same mutations were used in the CH2 and CH3 domain as for the full length bispecific IgG2a based TCE. In addition, the CH1 and hinge domains of both heavy chains were removed and replaced by a modified hinge, resulting in the following configuration: Chain A: VL_2C11_-GGGSGGG-VH_target_-GECPECP-CH2-CH3; Chain B: VL_target_-GGGSGGG-VH_2C11_-GRCPRCP-CH2-CH3. The same diabody TCE design was used for the DLL3-specific TCEs. The DLL3-specific antibodies DLL3-4, hSC16.56, and SC16.8 were used as templates for DLL3 TCEs.

### Recombinant protein expression and purification

All recombinant proteins were expressed in mammalian cells. Wild-type IL2 and IL2_REH_ were fused at the N-terminus to murine serum albumin (MSA) to extend *in vivo* half-life. Constructs for TRP2 full length IgG2a TCE, TRP2 diabody TCE, DLL3 diabody TCEs, murine wild-type MSA-IL2, MSA-IL2_REH_, and the sequence for wild-type murine interferon γ (mIFNγ) were cloned into the vector pD649 and transfected into Expi293 cells according to the manufacturer’s instructions. Briefly, Expi293 cells were transfected at a cell density of 3 x 10^6^ cells / mL using 1 µg of total DNA per mL cell culture. For bispecific IgG2a TCE four plasmids (2 different heavy chains and 2 different light chains) were used at 0.25 µg / mL culture volume, for bispecific diabody TCE two plasmids were cotransfected at 0.5 µg DNA / mL culture volume. Transfection enhancers were added ∼18 hours after transfection. Protein supernatant was harvested by centrifugation at 2500 g at 4 °C for 10 min on day 4 (mIFNγ, MSA-IL2, MSA-IL2_REH_) or day 5 (TCEs) after transfection. Cleared supernatant was filtered through a 0.22 µm filter. Large-scale production of diabody TCEs was performed in cell lines stably transduced to express the two diabody chains (ExpiCHO for TRP2 TCE, Expi293 for DLL3 TCE).

MSA-IL2, MSA-IL2_REH_, and mIFNγ were purified by standard immobilized metal ion affinity chromatography. PBS + 50 mM Tris-Cl was added at 1.5 x the supernatant’s volume, followed by the addition of Pierce High Capacity Ni-IMAC Resin and incubation at 4 °C overnight under constant agitation. Resin was washed 3 times with ∼ 10 resin volumes of HEPES-buffered saline (HBS) + 10 mM imidazole and proteins were eluted with HBS + 500 mM imidazole. TCEs were purified using Protein A affinity chromatography. Cleared and filtered supernatants were diluted with an equal volume of HBS and buffered by addition of Tris-Cl to a final concentration of 20 mM. Alkali-tolerant protein A agarose (MabSelect PrismA, Cytiva) was added, followed by incubation overnight at 4 °C under constant agitation. Protein A resin was washed 3 x times with ∼ 10 resin volumes of HBS + 20 mM Tris-Cl and eluted with 200 mM glycine (pH 2.5 – 3) directly into 2 M Tris-Cl for immediate pH-neutralization.

After concentration, recombinant proteins were polished by size-exclusion chromatography (SEC) on a Superdex® 200 Increase 10/300 GL (small scale preparations) or HiLoad® 16/600 Superdex® 200 pg (large scale preparations) on an ÄKTA pure™ chromatography system.

### Antigen density determination on tumor cell lines

B16F10 were either left untreated or incubated with 100 ng/mL IFNγ and/or 1 µM TRP2 peptide (SVYDFFVWL) for 24 hours, washed and stained with 100 nM TRP2 TCE or no antibody for 30 min in PBS followed by secondary staining with PE labelled anti-mouse IgG2a antibody (1:100, Thermo Fisher) antibody. The same staining procedure was used for the MC38 cell lines. The difference in geometric mean fluorescence intensity (ΔMFI) between identical cell lines stained with or without TRP2 TCE was plotted. The antibody bound per cell (ABC) for the SCLC cell lines was determined for anti-DLL3 PE (clone RMD3-13, BioLegend) using saturating concentrations of antibody and the BD Biosciences QuantiBRITE system.

### *In vitro* T cell activation and cytotoxicity assays

Splenocytes were prepared from C57BL/6 or OT-I mice by meshing through a 45 µm cell strainer followed by ammonium-chloride-potassium (ACK) red blood cell lysis. Splenocytes were either used directly or, for experiments involving effector T cells, first activated by plating cells onto 12-well plates precoated with 2.5 µg / mL anti-mCD3 (Clone 145-2C11, BioLegend) in the presence of 5 µg / mL anti-mCD28 (Clone 37.51, BioLegend) and 100 U / mL recombinant human IL2 (PeproTech). T cells were cultured for 5 to 7 days in 100 U / mL IL2, followed by a resting period of at least 24 hours in 20 U / mL IL2 before they were used in further experiments.

Following coculture of T cell and target cell lines for ∼36 hours, remaining target cells were briefly trypsinized, followed by staining with a cocktail containing 1:100 dilutions of the antibodies CD137-APC, CD8-BV785, CD69-PE for 15 min. Cells were washed in PBS and resuspended in PBS, containing SYTOX blue viability dye (1:1000). For SCLC cell lines (KP1, 16T, N2N1G) cells were stained without trypsinization in PBS containing 1:100 dilutions of the antibodies TCRβ-Percp/Cy5.5, DAPI, CD25-BV605, CD137-APC, CD8-BV785, CD69-PE for 15 min., followed by washing and resuspension in PBS containing DAPI (1:10 000). Cells were analyzed on a CytoFLEX-S (Beckman Coulter). Tumor cells were identified either by their lack of T cell marker expression (KP1, 16T, N2N1G) or expression of the transduced fluorescence markers eGFP (MC38 cell lines) or RFP (B16F10). The fraction of living target cells (SYTOX blue negative) out of all target cells was measured. Specific cytotoxicity was calculated as follows: *Specific lysis* = (1 − *targetviability* / max (*targetViability*)) ∗ 100.

### *In vitro* cytokine secretion assay

80,000 OT-I splenocytes were loaded with either 0.13 nM TRP2 (SVYDFFVWL) or Ova N4 (SIINFEKL) and either no TCE or 1 nM TRP2 Db TCE. Monensin (BioLegend) was added after 1 hour incubation at 37 °C, 5% CO2. After 12-hour incubation at 37 °C, 5% CO2 cells were spun down and labelled with TruStain FcX™ (1:100 in PBS; BioLegend). After 5 min incubation on ice, cells were stained with final concentrations of Zombie Aqua viability dye (1:1000, BioLegend), CD69-PE (1:100, BioLegend), CD3-Pacific Blue (1:100) for 15 mins on ice. Cells were then fixed using a Cytofix/Cytoperm™ Fixation/Permeabilization Kit (BD) according to the manufacturer’s instructions and stained with an antibody cocktail containing IL2-FITC (1:100, BioLegend), TCRβ-Percp/Cy5.5, CD4-APC/Cy7, CD8-BV785, IFNγ-PE/Cy7 (1:100) for 30 min on ice. Cells were washed, resuspended in PBS and analyzed on a CytoFLEX-S (Beckman Coulter).

### *In vitro* IFNγ treatment

5 x 10^5^ KP1 cells were seeded on day 0 with either 0nM or 1nM of murine IFNγ in RPMI with 10% Bovine Growth Serum and 1% penicillin–streptomycin and glutamine. One day later (day 1), cells were spun down, washed once with PBS, and switched to media without IFNγ. On day 2, cells were spun down, washed once with PBS, and kept on ice. Cells were then dissociated using 200 µL of Accutase. 1 mL of PBS was added to the cells in Accutase, and the cells were spun down. Cells were washed once with 200 µL of Annexin V binding buffer and then stained with FITC anti-mouse H-2 antibody (1:200 dilution from stock) in Annexin V binding buffer in the dark and on ice for 30 minutes. After staining, cells were washed once and resuspended in buffer containing DAPI (100 ng/mL). The cells were analyzed on a LSRFortessa Analyzer (BD). All spins were conducted at 400 g for 5 minutes.

### Tissue sample and flow cytometry

Tumor-draining lymph nodes and spleens were surgically removed, and single cell suspensions were generated by mincing through a 70 µm cell strainer. For spleens, RBC lysis was performed in 2 ml ACK for 2 min. at room temperature. Tumors were surgically removed and minced into small fragments using scissors. For flow cytometric analysis B16F10 were meshed through a 70 µm cell strainer and washed twice in PBS filtered through a 70 µm cell strainer and used for staining. KP1 tumors were minced and transferred to a 50 mL conical tube containing 9 mL of L15 media plus 0.25 mL of concentrated 5E enzyme mix and digested for 10 to 15 minutes, shaking at 37°C. After digestion, cells were filtered through a 70 µm cell strainer, spun down for 5 minutes at 400 g, washed with PBS, and RBC lysis was performed (using eBioscience™ 1X RBC Lysis Buffer). The concentrated 5E enzyme mix contains 4.25 mg/mL Collagenase I, 1.4 mg/mL Collagenase II, 4.25 mg/mL Collagenase IV, 0.625 mg/mL Elastase, and 0.625 mg/mL DNAse I in L15 medium. Cells were labeled with TruStain FcX™ (1:100 in PBS; BioLegend) followed by staining with the following monoclonal antibodies (1:100 in PBS unless otherwise specified) for B16F10: CD8-FITC, TRP2/K^b^ Tetramer-APC, Sca1-AF700, CD4-APC/Cy7, CD3-Pacific Blue, TCRγδ-BV605, CD11c-PE/Dazzle 594, TCRβ-Percp/Cy5.5, CD45.2-PE/Cy7; for KP1: CD11b-AF488, TCRβ-Percp/Cy5.5, CD25-APC, CD45.2-AF700, CD4-APC/Cy7, CD3-Pacific Blue, Zombie Aqua (1:1000), CD8-BV785, I-Ab-PE, CD11c-PE/Dazzle 594; and CD44-FITC, TCRβ-Percp/Cy5.5, PD1-APC, CD45.2-AF700, CD62L-APC/Cy7, CD3-Pacific Blue, Zombie Aqua (1:1000), Lag3-BV650, IL7RA-BV785, TIM3-PE/Cy5, TIGIT-PE/Cy7, CD8-PE; and H2-K^b^-APC, CD45.2-AF700, DLL3-PE, Zombie Aqua (1:1000). Cells were stained for 30 min. on ice, washed with PBS and analyzed on a CytoFLEX-S (Beckman Coulter).

### Single-cell genomics sample preparation

On day 6 after treatment with HBS, IL2_REH_ alone, TCE alone, or combination TCE and IL2_REH_, B16F10 tumors were surgically removed and dissociated using a custom protocol to remove debris and enrich for live TILs. Specifically, 7-8 tumors from each treatment group were dissociated using a mouse Tumor Dissociation Kit (Miltenyi) on a GentleMACS Octo Dissociator using program 37C_m_TDK_1. Single cell suspensions were filtered through a cell strainer, pooled by treatment group and lymphocytes were isolated by Ficoll density gradient centrifugation, followed by T cell enrichment with mouse CD4/CD8 (TIL) MicroBeads (Miltenyi). Enriched TILs were then labelled in PBS with lipid-modified oligonucleotides (LMOs) hybridized to a sample-specific MULTI-seq barcode as described previously (*36*). MULTI-seq LMO labelling reactions were then quenched with 1% BSA in PBS, pooled, and washed with 1% BSA in PBS prior to labelling with TruStain FcX™ (1:100 in PBS; BioLegend). After 5 minutes on ice, 25 µL of antibody cocktail was added to achieve final staining concentrations of 1:100 anti-CD8-FITC (FITC; BD), 1:100 anti-murine TCR β chain (PE; BioLegend), Zombie NIR viability dye (1:500, BioLegend). After 30 minutes on ice, the cells were diluted with 1 mL of FACS buffer, washed once with 1 mL of FACS buffer, and filtered through a 70µm Macs SmartStrainer prior to FACS enrichment for live mTCR β chain^+^ cells using a BD FACSAria III instrument. After FACS, cells were counted, the concentration was adjusted to 1 x 10^6^ cells/mL, and the cell suspension was ‘super-loaded’ across 3 lanes of a 10x Genomics 5’ scRNA-seq V2 microfluidics chip.

### Single-cell genomics library preparation and next generation sequencing

scRNA-seq and scTCR-seq libraries were prepared according to manufacturer’s recommendations (10x Genomics; CG000331 Rev F). MULTI-seq libraries were prepared as described previously (*36*). scRNA-seq, scTCR-seq, and MULTI-seq libraries were pooled and sequenced using the Illumina NovaSeqX platform (25B flow cell). scRNA-seq, TCR-seq, and MULTI-seq libraries were sequenced to an average of 47,851, 14,753, and 1,872 reads per cell, respectively.

### Single-cell genomics data pre-processing and quality-control

scRNA-seq and scTCR-seq library FASTQs were pre-processed using Cell Ranger version 7.0.0 (10x Genomics) and aligned to the mm-10-3.0.0 reference transcriptome and GRCm38 V(D)J reference, respectively. Cell Ranger aggregate was used to perform scTCR-seq read-depth normalization and clonotype aggregation. Filtered scRNA-seq count matrices were then read into R and parsed to exclude genes with fewer than 5 counts across all cell barcodes. Parsed scRNA-seq data was then pre-processed using Seurat V5 (*53*), and clusters with low total UMIs and/or high proportion of mitochondrial transcripts were excluded. Cell barcodes passing the first quality-control workflow were then used to pre-process MULTI-seq barcode FASTQs and perform sample classification using the ‘deMULTIplex2’ R package (*54*) with the additional semi-supervised negative cell rescue workflow from the ‘deMULTIplex’ R package (*36*). Following MULTI-seq demultiplexing, unclassified cells and clusters enriched with MULTI-seq-defined doublets were removed prior to re-processing, as described previously (*36*). These data were used for unsupervised clustering, differential gene expression testing, and manual annotation of major TIL subtypes, as described in the Results.

After the initial scRNA-seq and MULTI-seq analyses, filtered scTCR-seq data was read into R and parsed to only include cells passing the scRNA-seq and MULTI-seq quality-control workflows. The scTCR-seq data was further parsed to remove cells with biologically impossible TCR chain usage, and a subsetted data object containing high-quality paired scRNA-seq and scTCR-seq data was computed using Seurat. These data were used for comparing patterns of T cell clonal diversity between TIL subtypes and treatment groups.

### Statistical analysis

Longitudinal analysis of tumor sizes was carried out by linear mixed-effect modeling of log-transformed tumor volumes using the R package TumGrowth (*55*). For graphing, the mean of the untransformed tumor volumes of each group ± standard error of the mean are shown. Type II ANOVA was performed to test if tumor growth-rates differ between groups. P-values were adjusted according to the Holm method for all pairwise comparisons of interest. Statistical comparison of cell numbers between groups was performed using the stat_pwc function of the ggpubr R package. All main text and supplemental figures were generated using the ggplot2, ggpubr, scales, and paletteer R packages.

Statistically-significant shifts in TIL subtype proportions in the scRNA-seq data were identified using the ‘propeller’ function with bootstrapping in the ‘Speckle’ R package (*56*). Differentially-expressed genes between clusters in all datasets were defined using the Wilcoxon rank-sum test as implemented in the ‘FindAllMarkers’ Seurat function (log fold-change thresholds used varied between 2-2.5 and minimum proportion expressed varied between 0.5-0.8, depending on number of desired features in each visualization). Specific differential expression testing was performed using the Wilcoxon rank-sum test as implemented in the ‘FindMarkers’ Seurat function. All main text and supplemental figures describing single-cell genomics analyses were generated using the Seurat, ggplot2, and ComplexHeatmap R packages (*57*).

## Supplementary Materials

Supplemental Figure S1: *In vitro* development of TCEs, related to Fig. 1.

Supplemental Figure S2: TCE monotherapy against tumors with low and high antigen density, related to Fig. 2.

Supplemental Figure S3: TCE IL2-combination therapy against low antigen-density tumors, related to Fig. 3.

Supplemental Figure S4, TCE + IL2 combination induced toxicity, related to Fig. 3.

Supplemental Figure S5: Multimodal single-cell analysis of tumor infiltrating lymphocytes across treatment groups, related to Fig. 4.

Supplemental Table S1: **A**mino acid sequences of the TCEs used in the current study.

## Acknowledgments

We thank Zev Gartner (UCSF) for providing MULTI-seq reagents and Antoni Ribas (UCLA) for providing cell lines. Cell sorting for this project was done in part on instruments in the Stanford Shared FACS Facility (RRID: SCR_017788). Sequencing was performed at the UCSF Center for Advanced Technology. We thank Christopher Chang and Chang Liu (UCSF) for collaboration related to TRP2 TCEs.

## Funding

National Institutes of Health grant 3U54CA24471101 (KCG)

National Institutes of Health grant RO1AI-51321 (KCG)

Parker Institute of Cancer Immunotherapy (KCG, JS, ATS, RAHL, JE)

Yosemite Innovation Fund (KCG, ATS)

Mark Foundation Endeavor Grant (KCG, JS, RAHL)

Mark Foundation Emerging Leader Award (ATS)

Berlin Institute of Health Charité Clinician Scientist Program (MO)

METAVivor Early Career Investigator Award (CSM)

National Institutes of Health grant 1K99CA293137-01A1 (CSM)

Pew-Stewart Scholars for Cancer Research Award (ATS)

Cancer Research Institute Lloyd J. Old STAR Award (ATS)

Burroughs Wellcome Fund Career Award at the Scientific Interface (RAHL)

Biohub San Francisco (RAHL)

Howard Hughes Medical Institute (KCG)

Cancer Research Institute Irvington Postdoctoral Fellowship (JM)

CRISPR Cures for Cancer (JE)

Grand Multiple Myeloma Translational Initiative (JE)

Schweizerischer Nationalfonds (SNSF)

Postdoctoral Fellowship (CP)

Deutsche Forschungsgemeinschaft (DFG)

Walter Benjamin-Program grant CH2718/1-1 (XC)

## Author contributions

MO conceived and designed the study with input from KCG, JS, and ATS. CSM, CLP, and MO performed data analysis. MO, CLP, CSM, CP, WY, HJ, XJ, JJM, LLS, ZM, and DW performed experiments. ATS, JE, RAHL, JS, and KCG provided resources. MO, CLP, CSM, ATS, JS, and KCG wrote the manuscript with input from all authors.

## Data and materials availability

All reagents generated in this study are available from the lead contact (KCG) upon request and with a completed materials transfer agreement if needed. Single-cell RNA-seq data have been deposited at GEO (GSEXXXX) and are publicly available as of the date of publication. Any additional information required to reanalyze the data reported in this paper is available from the lead contact (KCG) upon request. Processed data and R scripts used for analyzing data and generating all manuscript figures are deposited on GitHub: github.com/chris-mcginnis-ucsf/tce_clonal_replacement.

