## Supplementary materials for "T cell engagers control solid tumors through clonal replacement and IL2-driven effector differentiation of CD8 T cells"

### Table S1 amino acid sequences of the TCEs used in the current study

| | | |
|---|---|---|
| TRP2 IgG2 TCE | HA | EVLLQQSGPELVKPGASVKIPCKASGYTFTDYYMDWVKQSHGKSLEWIGLINPNNGGTV<br>YNQKFKGKATLTVDKSSSVAYMEVRSLTSEDTAVYYCARKPYTGHPAWFAYWGQGTLV<br>TVSAAKTTAPSVYPLAPVCGDTTGSSVTLGCLVKGYFPEPVTLTWNSGSLSSGVHTFPA<br>VLQSDLYTLSSSVTVTSSTWPSQSITCNVAHPASSTKVDKKIEPRGPTIKPCPPCKCPAPN<br>LLGGPSVFIFPPKIKDVLMISLSPIVTCVVVAVSEDDPDVQISWFVNNVEVHTAQTQTHRE<br>DYNSTLRVVSALPIQHQDWMSGKEFKCKVNNKDLPAPIERTISKPKGSVRAPQVYVLPPP<br>EEEMTKKQVSLTCLVKDFMPEDIYVEWTNNGKTELNYKNTEPVLDSDGSYFMYSELTVE<br>KKNWVERNSYSCSVVHEGLHNHHTTDSFSRTPGK |
| | HB | EVQLVESGGGLVQPGKSLKLSCEASGFTFSGYGMHWVRQAPGRGLESVAYITSSSINIK<br>YADAVKGRFTVSRDNAKNLLFLQMNILKSEDTAMYYCARFDWDKNYWGQGTMVTVSSA<br>KTTAPSVCPLAPVVGDTTGSSVTLGCLVKGYFPEPVTLTWNSGSLSSGVHTFPAVLQSD<br>LYTLSSSVTVTSSTWPSQSITCNVAHPASSTKVDKKIEPRGPTIKPCPPCKCPAPNLLGGP<br>SVFIFPPKIKDVLMISLSPIVTCVVVAVSEDDPDVQISWFVNNVEVHTAQTQTHREDYNST<br>LRVVSALPIQHQDWMSGKEFKCKVNNKDLPAPIERTISKPKGSVRAPQVYVLPPPEKEMT<br>KKQVSLTCLVKDFMPEDIYVEWTNNGKTELNYKNTEPVLKSDGSYFMYSKLTVEKKNWV<br>ERNSYSCSVVHEGLHNHHTTKSFSRTPGK |
| | LC A | DIQVTQSSSSFSVSLGDRVTITCKASEDIYNRLAWYQQKPGNAPRLLISGATSLETGVPDR<br>FSGSGSRKDYTLIITSLQTEDVATYYCQQYYAFPLTFGAGTKLELKRADAAPTVSIFPPSS<br>EQLTSGGASVVCFLNNFYPKDINVKWKIDGSERQNGVLNSWTDQDSKDSTYSMSSTLTL<br>TKDEYERHNSYTCEATHKTSTSPIVKSFNRNEC |
| | LC B | DIQMTQSPSSLPASLGDRVTINCQASQDISNYLNWYQQKPGKAPKLLIYYTNKLADGVPS<br>RFSGSGSGRDSSFTISSLESEDIGSYYCQQYYNYPWTFGPGTKLEIKRADAAPTVSIFPP<br>CSEQLTSGGASVVCFLNNFYPKDINVKWKIDGSERQNGVLNSWTDQDSKDSTYSMSSTL<br>TLTKDEYERHNSYTCEATHKTSTSPIVKSFNRNEV |
| TRP2 Db TCE | A | DIQMTQSPSSLPASLGDRVTINCQASQDISNYLNWYQQKPGKAPKLLIYYTNKLADGVPS<br>RFSGSGSGRDSSFTISSLESEDIGSYYCQQYYNYPWTFGPGTKLEIKGGGSGGGEVLLQ<br>QSGPELVKPGASVKIPCKASGYTFTDYYMDWVKQSHGKSLEWIGLINPNNGGTVYNQKF<br>KGKATLTVDKSSSVAYMEVRSLTSEDTAVYYCARKPYTGHPAWFAYWGQGTLVTVSAG<br>ECPECPAPNLLGGPSVFIFPPKIKDVLMISLSPIVTCVVVAVSEDDPDVQISWFVNNVEVH<br>TAQTQTHREDYNSTLRVVSALPIQHQDWMSGKEFKCKVNNKDLPAPIERTISKPKGSVRA<br>PQVYVLPPPEEEMTKKQVSLTCLVKDFMPEDIYVEWTNNGKTELNYKNTEPVLDSDGSY<br>FMYSELTVEKKNWVERNSYSCSVVHEGLHNHHTTDSFSRTPGK |
| | B | DIQVTQSSSSFSVSLGDRVTITCKASEDIYNRLAWYQQKPGNAPRLLISGATSLETGVPDR<br>FSGSGSRKDYTLIITSLQTEDVATYYCQQYYAFPLTFGAGTKLELKGGGSGGGEVQLVES<br>GGGLVQPGKSLKLSCEASGFTFSGYGMHWVRQAPGRGLESVAYITSSSINIKYADAVKG<br>RFTVSRDNAKNLLFLQMNILKSEDTAMYYCARFDWDKNYWGQGTMVTVSSGRCPRCPA<br>PNLLGGPSVFIFPPKIKDVLMISLSPIVTCVVVAVSEDDPDVQISWFVNNVEVHTAQTQTH<br>REDYNSTLRVVSALPIQHQDWMSGKEFKCKVNNKDLPAPIERTISKPKGSVRAPQVYVLP<br>PPEKEMTKKQVSLTCLVKDFMPEDIYVEWTNNGKTELNYKNTEPVLKSDGSYFMYSKLT<br>VEKKNWVERNSYSCSVVHEGLHNHHTTKSFSRTPGK |
| DLL3 Db TCE (DLL3-4) | A | DIQMTQSPSSLPASLGDRVTINCQASQDISNYLNWYQQKPGKAPKLLIYYTNKLADGVPS<br>RFSGSGSGRDSSFTISSLESEDIGSYYCQQYYNYPWTFGPGTKLEIKGGGSGGGQVQLQ<br>ESGPGLVKPSETLSLTCTVSGGSISSYYWSWIRQPPGKGLEWIGYVYYSGTTNYNPSLK<br>SRVTISVDTSKNQFSLKLSSVTAADTAVYYCASIAVTGFYFDYWGQGTLVTVSSGECPEC<br>PAPNLLGGPSVFIFPPKIKDVLMISLSPIVTCVVVAVSEDDPDVQISWFVNNVEVHTAQTQ<br>THREDYNSTLRVVSALPIQHQDWMSGKEFKCKVNNKDLPAPIERTISKPKGSVRAPQVYV<br>LPPPEEEMTKKQVSLTCLVKDFMPEDIYVEWTNNGKTELNYKNTEPVLDSDGSYFMYSE<br>LTVEKKNWVERNSYSCSVVHEGLHNHHTTDSFSRTPGK |
| | B | EIVLTQSPGTLSLSPGERVTLSCRASQRVNNNYLAWYQQRPGQAPRLLIYGASSRATGIP<br>DRFSGSGSGTDFTLTISRLEPEDFAVYYCQQYDRSPLTFGGGTKLEIKGGGSGGGEVQL<br>VESGGGLVQPGKSLKLSCEASGFTFSGYGMHWVRQAPGRGLESVAYITSSSINIKYADA<br>VKGRFTVSRDNAKNLLFLQMNILKSEDTAMYYCARFDWDKNYWGQGTMVTVSSGRCPR<br>CPAPNLLGGPSVFIFPPKIKDVLMISLSPIVTCVVVAVSEDDPDVQISWFVNNVEVHTAQT<br>QTHREDYNSTLRVVSALPIQHQDWMSGKEFKCKVNNKDLPAPIERTISKPKGSVRAPQV |

**Table S1 amino acid sequences of the TCEs used in the current study**

| | | |
|---|---|---|
| DLL3 Db TCE (hSC16.56) | A | YVLPPPEKEMTKKQVSLTCLVKDFMPEDIYVEWTNNGKTELNYKNTEPVLKSDGSYFMY<br>SKLTVEKKNWVERNSYSCSVVHEGLHNHHTTKSFSRTPGK<br>DIQMTQSPSSLPASLGDRVTINCQASQDISNYLNWYQQKPGKAPKLLIYYTNKLADGVPS<br>RFSGSGSGRDSSFTISSLESEDIGSYYCQQYYNYPWTFGPGTKLEIKGGGSGGGQVQLV<br>QSGAEVKKPGASVKVSCKASGYTFTNYGMNWVRQAPGQGLEWMGWINTYTGEPTYAD<br>DFKGRVTMTTDTSTSTAYMELRSLRSDDTAVYYCARIGDSSPSDYWGQGTLVTVSSGEC<br>PECPAPNLLGGPSVFIFPPKIKDVLMISLSPIVTCVVVAVSEDDPDVQISWFVNNVEVHTA<br>QTQTHREDYNSTLRVVSALPIQHQDWMSGKEFKCKVNNKDLPAPIERTISKPKGSVRAP<br>QVYVLPPPEEEMTKKQVSLTCLVKDFMPEDIYVEWTNNGKTELNYKNTEPVLDSDGSYF<br>MYSELTVEKKNWVERNSYSCSVVHEGLHNHHTTDSFSRTPGK |
| | B | EIVMTQSPATLSVSPGERATLSCKASQSVSNDVVWYQQKPGQAPRLLIYYASNRYTGIPA<br>RFSGSGSGTEFTLTISSLQSEDFAVYYCQQDYTSPWTFGQGTKLEIKRGGGSGGGEVQL<br>VESGGGLVQPGKSLKLSCEASGFTFSGYGMHWVRQAPGRGLESVAYITSSSINIKYADA<br>VKGRFTVSRDNAKNLLFLQMNILKSEDTAMYYCARFDWDKNYWGQGTMVTVSSGRCPR<br>CPAPNLLGGPSVFIFPPKIKDVLMISLSPIVTCVVVAVSEDDPDVQISWFVNNVEVHTAQT<br>QTHREDYNSTLRVVSALPIQHQDWMSGKEFKCKVNNKDLPAPIERTISKPKGSVRAPQV<br>YVLPPPEKEMTKKQVSLTCLVKDFMPEDIYVEWTNNGKTELNYKNTEPVLKSDGSYFMY<br>SKLTVEKKNWVERNSYSCSVVHEGLHNHHTTKSFSRTPGK |
| DLL3 Db TCE (SC16.8) | A | DIQMTQSPSSLPASLGDRVTINCQASQDISNYLNWYQQKPGKAPKLLIYYTNKLADGVPS<br>RFSGSGSGRDSSFTISSLESEDIGSYYCQQYYNYPWTFGPGTKLEIKGGGSGGGEQAQL<br>QQSGAELVRPGTSVKVSCKASGYAFTNYLIEWVKQRPGQGLEWIGVINPGTGGTNYNEN<br>FKGKATLTADKSSSTAYMQLSSLTSDDSAVYFCARSPYDYHEGAMDYWGQGTSVTVSS<br>GECPECPAPNLLGGPSVFIFPPKIKDVLMISLSPIVTCVVVAVSEDDPDVQISWFVNNVEV<br>HTAQTQTHREDYNSTLRVVSALPIQHQDWMSGKEFKCKVNNKDLPAPIERTISKPKGSV<br>RAPQVYVLPPPEEEMTKKQVSLTCLVKDFMPEDIYVEWTNNGKTELNYKNTEPVLDSDG<br>SYFMYSELTVEKKNWVERNSYSCSVVHEGLHNHHTTDSFSRTPGK |
| | B | EIQMTQSPSSMSASLGDRITITCQATQDIVKNLNWYQQKPGKPPSFLIYYAIELAEGVPSR<br>FSGSGSGSDYSLTISNLESEDFADYYCLQFYEFPFTFGAGTKLELKGGGSGGGEVQLVE<br>SGGGLVQPGKSLKLSCEASGFTFSGYGMHWVRQAPGRGLESVAYITSSSINIKYADAVK<br>GRFTVSRDNAKNLLFLQMNILKSEDTAMYYCARFDWDKNYWGQGTMVTVSSGRCPRC<br>PAPNLLGGPSVFIFPPKIKDVLMISLSPIVTCVVVAVSEDDPDVQISWFVNNVEVHTAQTQ<br>THREDYNSTLRVVSALPIQHQDWMSGKEFKCKVNNKDLPAPIERTISKPKGSVRAPQVYV<br>LPPPEKEMTKKQVSLTCLVKDFMPEDIYVEWTNNGKTELNYKNTEPVLKSDGSYFMYSK<br>LTVEKKNWVERNSYSCSVVHEGLHNHHTTKSFSRTPGK |

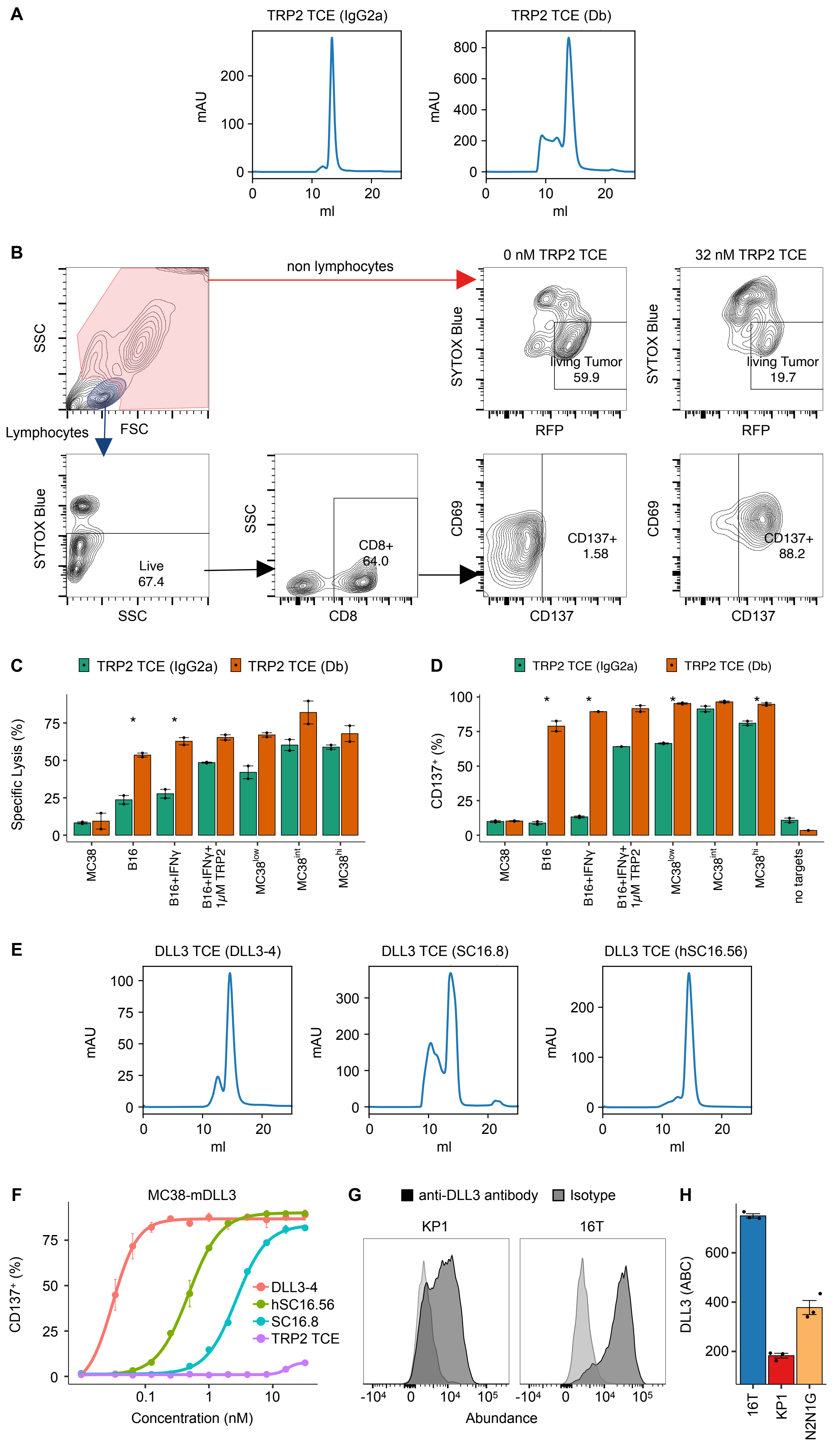


**Figure S1: I*n vitro* development of TCEs, related to Fig. 1, legend on next page.**

**Figure S1: I*n vitro* development of TCEs, related to Fig. 1**

(A) Representative size-exclusion chromatography traces for TRP2 IgG2a TCE (left) and TRP2 Db TCE (right) on a Superdex 200 Increase, 10/300 GL column.

(B) Gating-strategy to identify T cells and target cells during co-culture to determine target cell viability and CD137 expression on CD8+ T cells.

(C) Comparison of cytotoxicity (specific lysis) induced by TRP2 IgG2a TCE (green) and TRP2 Db TCE (orange) at a TCE concentration of 32 nM; values represent mean ±s.e.m. of intra-assay duplicates; *: $p < 0.05$ (student's t-test).

(D) Comparison of CD137 up-regulation induced by TRP2 IgG2a TCE (green) and TRP2 Db TCE (orange) at a TCE concentration of 32 nM; values represent mean ± s.e.m. of intra-assay duplicates; *: $p < 0.05$ (student's t-test).

(E) Representative size-exclusion chromatography traces for DLL3 Db TCEs with 3 different DLL3 binders (left: DLL3-4, middle: SC16.8, right: hSC16.56) on a Superdex 200 Increase, 10/300 GL column.

(F) MC38 over-expressing murine DLL3 were co-cultured with mouse effector T cells at an Effector-to-target-ratio of 1:1 (40 000 target cells per well) in the presence of the indicated concentrations of 3 different DLL3 specific TCEs or TRP2 TCE as control. T cell activation was assessed by staining for CD137 surface expression on CD8+ T cells after 36 hours. Data represent mean ± s.e.m. of intra-assay duplicates.

(G) KP1 (left) and 16T SCLC (right) were stained with an anti-DLL3 antibody or isotype control to compare DLL3 expression.

(H) Comparison of DLL3 expression across SCLC stained with anti-DLL3 antibody and calibrated with BD Quantibrite beads.

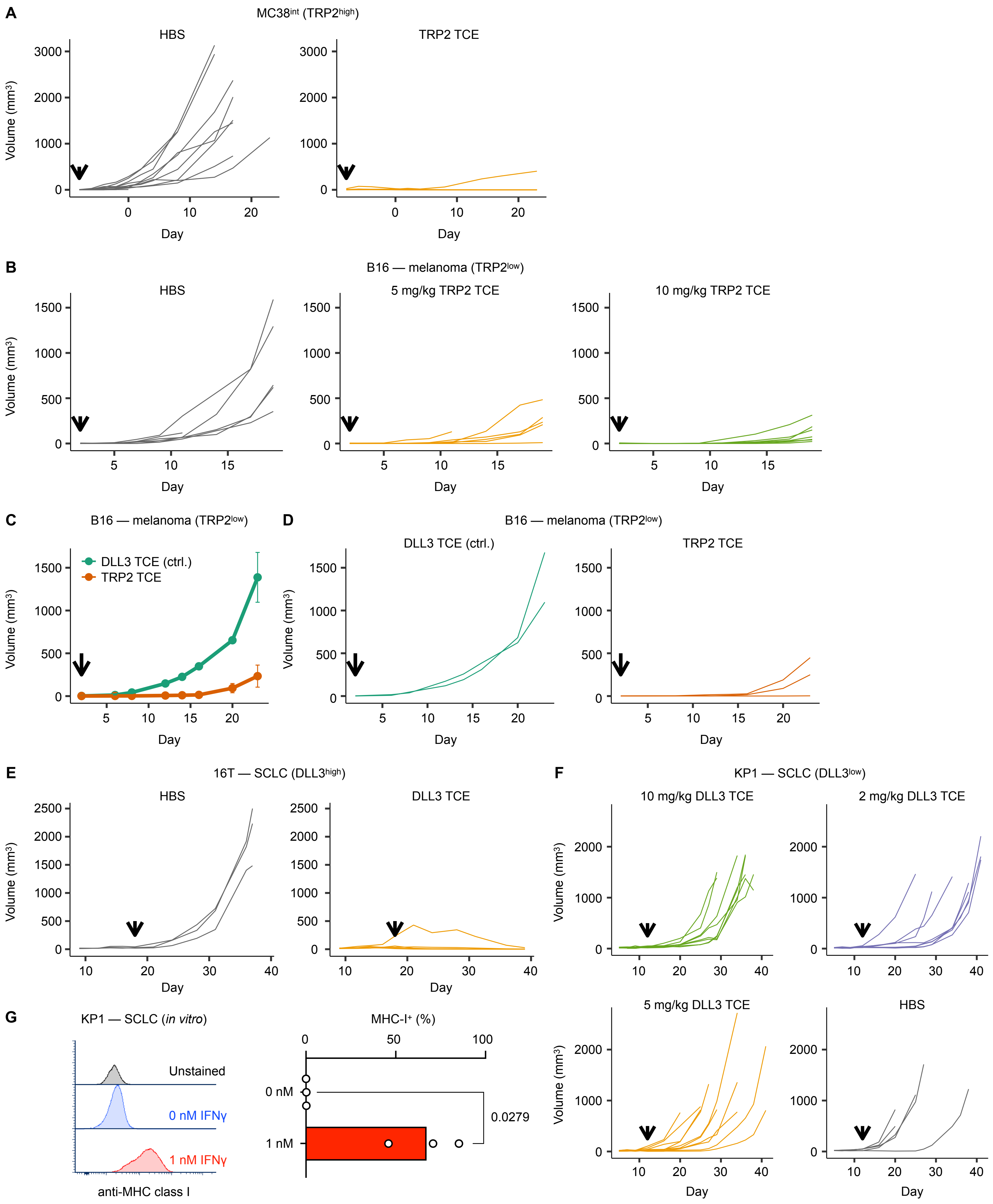


**Figure S2: TCE monotherapy against tumors with low and high antigen density, related to Fig. 2, legend on next page**

**Figure S2: TCE monotherapy against tumors with low and high antigen density, related to Fig. 2**

(A) Tumor growth curves of individual mice bearing MC38-Trp2 tumors treated with control or TRP2 TCE. Arrows indicate treatment start.

(B) Tumor growth curves of individual B16F10 tumor bearing mice treated with HBS (left), 5 mg/kg TRP2 TCE (middle), or 10 mg/kg TRP2 TCE (right). Arrows indicate treatment start.

(C) B16F10 tumor bearing mice were treated with 5 mg / kg DLL3 TCE (control, green, middle) or 5 mg/kg TRP2 TCE (orange, right). Left: values represent mean ± s.e.m. Tumor growth between groups was not significantly different (Linear mixed-effects modeling). Arrows indicate treatment start.

(D) Individual tumor growth curves from experiment in (C).

(E) Tumor growth curves for individual mice bearing 16T ($DLL3_{high}$) tumors treated with HBS (left) or DLL3 TCE (right). Arrows indicate treatment start.

(F) Tumor growth curves for individual mice bearing KP1 ($DLL3_{low}$) tumors treated with 3 different dose levels of DLL3 TCE or HBS. Arrows indicate treatment start.

(G) KP1 SCLC cells were left untreated or incubated with 1 nM murine IFNγ for 24 hours, washed and stained for expression of MHC class I. Bar graphs show mean ± s.e.m. of three independent experiments

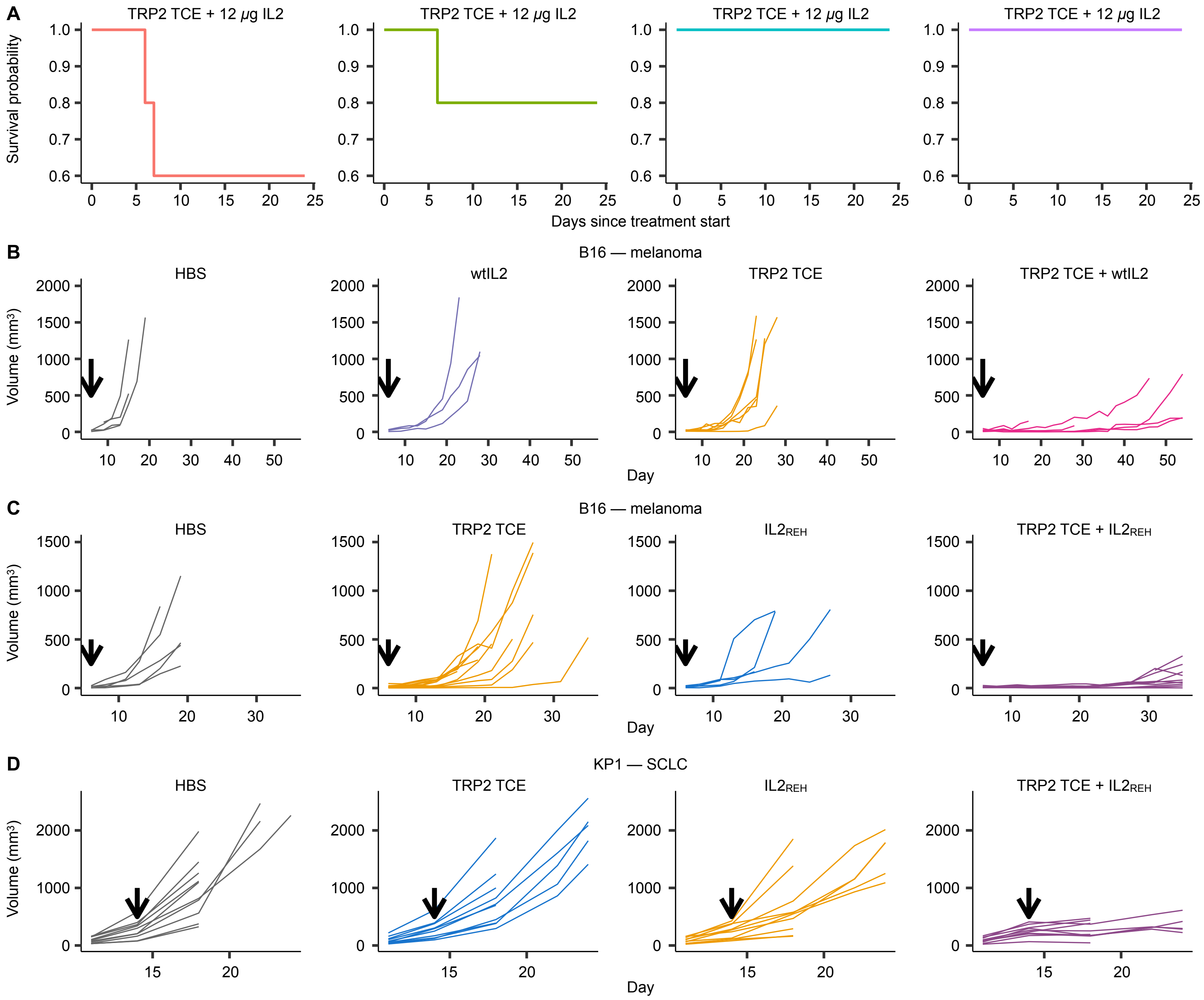


**Figure S3: TCE IL2-combination therapy against low antigen-density tumors, related to Fig. 3**

(A) Tumor-free C57BL/6 mice were treated with 5 mg/kg TRP2 TCE q6d and the indicated doses of wild-type murine IL2 q2d to determine tolerability of the IL2+TCE combination. Shown is the fraction of surviving animals since treatment start.

(B) Tumor growth curves for individual B16F10 melanoma bearing animals treated with HBS, wild-type IL2, TRP2 TCE or TRP2 TCE + wild-type IL2.

(C) Tumor growth curves for individual B16F10 melanoma bearing animals treated with HBS, $IL2_{REH}$, TRP2 TCE or TRP2 TCE + $IL2_{REH}$.

(D) Tumor growth curves for individual KP1 SCLC bearing animals treated with HBS, $IL2_{REH}$, DLL3 TCE or DLL3 TCE + $IL2_{REH}$.

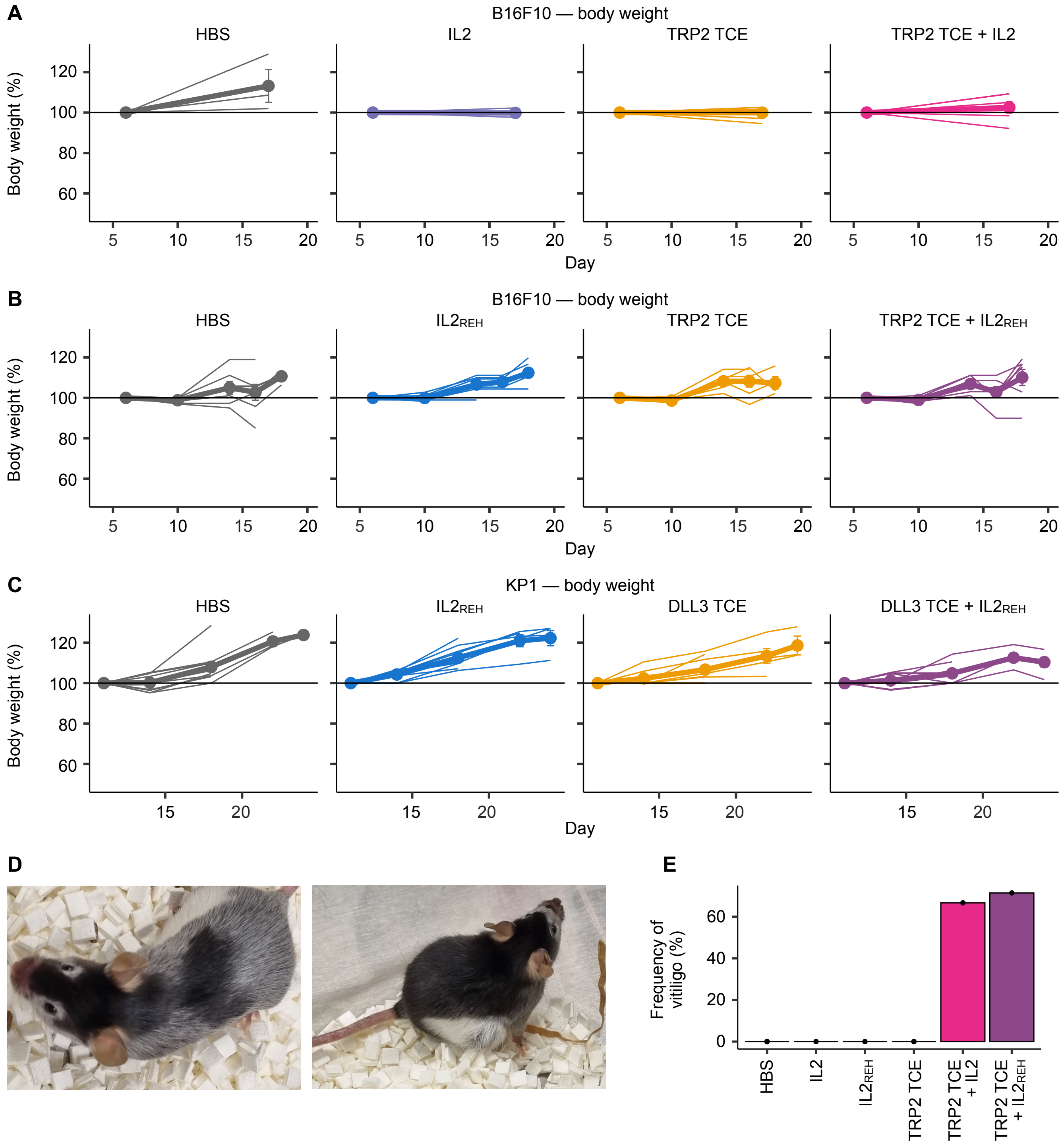


**Figure S4: TCE + IL2 combination induced toxicity, related to Fig. 3.**

(A) Body weight changes during treatment of B16F10 treated with TRP2 TCE and/or wtIL2.

(B) Body weight changes during treatment of B16F10 treated with TRP2 TCE and/or $IL2_{REH}$.

(C) Body weight changes during treatment of KP1 tumors treated with DLL3 TCE and/or $IL2_{REH}$.

(D) Examples of two animals under TRP2 TCE + IL2 developing on-target/off-tumor toxicity (leukotrichia, left, middle).

(E) Bar graphs showing the frequency of animals developing vitiligo/leukotrichia during treatment under treatment with IL2 / $IL2_{REH}$ + TRP2 TCE compared to other treatment groups.

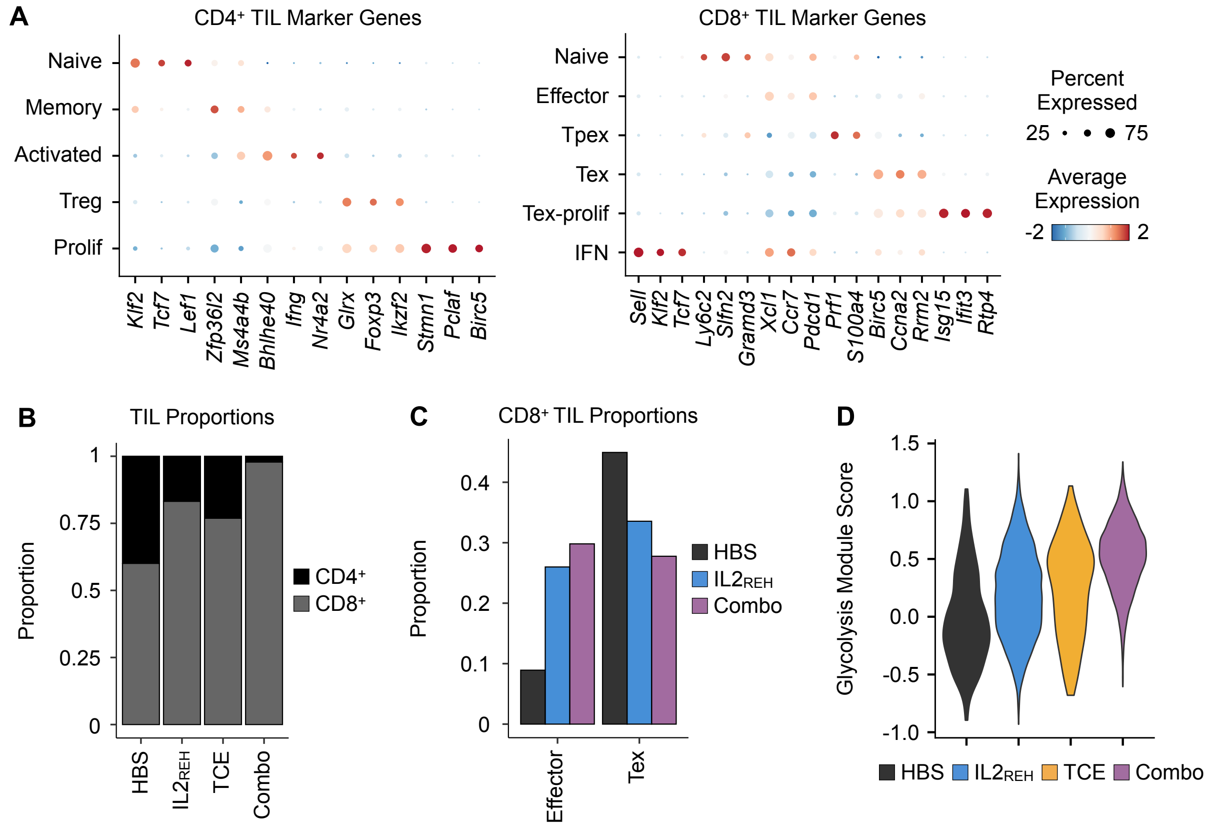


**Figure S5: Multimodal single-cell analysis of tumor infiltrating lymphocytes across treatment groups, related to Fig. 4.**

(A) Dot plot of $CD4^+$ TIL (left) and $CD8^+$ TIL (right) subtype annotation genes. Dot color indicates expression level and size indicates the proportion of cells expressing each gene.

(B) Bar chart of $CD4^+$ and $CD8^+$ TIL proportions in each treatment group.

(C) Bar chart of effector and Tex $CD8^+$ TIL proportions in each $IL2_{REH}$ treatment arm and controls.

(D) Violin plots of glycolysis gene signature module scores in effector $CD8^+$ TILs in each treatment group.
